# SERINC1-dependent lipid preservation gates neocortical synaptogenesis and remote memory

**DOI:** 10.64898/2026.09.15.751742

**Authors:** Shaohua Wu, Haijin Chen, Xiao Liu, Xiaoxia Lin, Xian Jiang

## Abstract

Systems memory consolidation reorganizes episodic memories from the hippocampus into neocortical networks such as the anterior cingulate cortex (ACC), yet the localized molecular programs supporting this process remain unknown. Here, we show that the transmembrane lipid regulator *Serinc1* is selectively induced in the ACC during neocortical memory consolidation. Loss of *Serinc1* reduces ACC synapse density, blocks learning-induced *de novo* callosal synaptogenesis, and selectively abolishes remote memory. Lipidomic profiling reveals that SERINC1 prevents activity-dependent depletion of inner-leaflet anionic phospholipids—specifically docosahexaenoic acid (DHA) bearing-phosphatidylserine and arachidonic acid (AA) bearing-phosphatidylinositol—safeguarding signaling cascades required for structural plasticity. Targeted viral restoration of SERINC1 in the ACC rescues cortical synapse density and remote memory. Thus, long-term memory is gated by a localized program of activity-dependent neuro-lipidomic remodeling.

## Main Text

The long-term storage of episodic experiences relies on systems memory consolidation, a process wherein memory traces initially dependent on the hippocampus are progressively reorganized into distributed neocortical networks, including the anterior cingulate cortex (ACC) (*1–6*). Although systems-level circuit reorganization during memory consolidation is well documented, the localized, cortex-specific molecular programs that enable neocortical circuits to establish and maintain newly formed synaptic connections remain elusive. Canonical signaling cascades, such as the cAMP–PKA–CREB axis, serve as ubiquitous mediators of plasticity across multiple memory phases and brain regions (*7–10*), yet do not fully explain the regionalized, temporally dynamic structural remodeling characteristic of neocortical hubs (*11–13*). We reasoned that a specialized transcriptional or metabolic program must be selectively recruited within neocortical structures during offline consolidation to remodel the local biophysical microenvironment and protect emerging circuits from structural destabilization.

Mining our single-cell RNA sequencing atlas of remote-memory-associated transcriptional programs in the medial prefrontal cortex (mPFC), we identified *Serinc1* (Serine Incorporator 1), encoding a transmembrane lipid regulator, as a key target broadly induced in mPFC during memory consolidation (*14*). SERINC family proteins facilitate serine incorporation into cellular phospholipids and modulate membrane symmetry (*15–17*), while membrane lipids broadly regulate neurotransmission and synaptic plasticity (*18–22*). However, whether neocortical lipids actively gate long-term memory stabilization—and specifically how SERINC1 coordinates distinct phospholipid microenvironments during systems memory consolidation—remains unknown.

Here, we investigated whether localized neocortical lipid regulation acts as an obligate gatekeeper for structural plasticity and long-range circuit assembly during memory consolidation. Combining region- and cell-type-specific knockouts, lipidomic and metabolomic profiling, dual-virus mGRASP imaging, and viral rescue, we show that SERINC1-dependent lipid preservation prevents activity-dependent collapse of anionic phospholipid pools to sustain neocortical synaptogenesis and remote memory.

## Results

### *Serinc1* is selectively induced in the ACC and required for remote memory

To determine whether *Serinc1* expression is dynamically regulated during systems memory consolidation, we mapped its spatiotemporal dynamics across hippocampal CA1 and layer II/III of the ACC—key nodes for recent memory retention and remote engram consolidation, respectively (*11, 13, 23*). Wild-type mice were subjected to contextual fear conditioning (CFC) paired with multiplexed fluorescence in situ hybridization (RNAscope) (Fig. 1, A and B). Fear-conditioned (FR) mice and unconditioned no-fear controls (NF) were tested for contextual memory retrieval at either recent (Day 1) or remote (Day 16) time points post-training. Following recent memory recall, *Serinc1* mRNA abundance remained unchanged in both the hippocampal CA1 and layer II/III of the ACC relative to NF controls (Fig. 1, C and D). In contrast, remote memory retrieval elicited a robust, region-specific upregulation of *Serinc1* transcripts in ACC layer II/III, whereas hippocampal CA1 *Serinc1* expression remained strictly locked at baseline (Fig. 1, E and F).

**Fig. 1.**
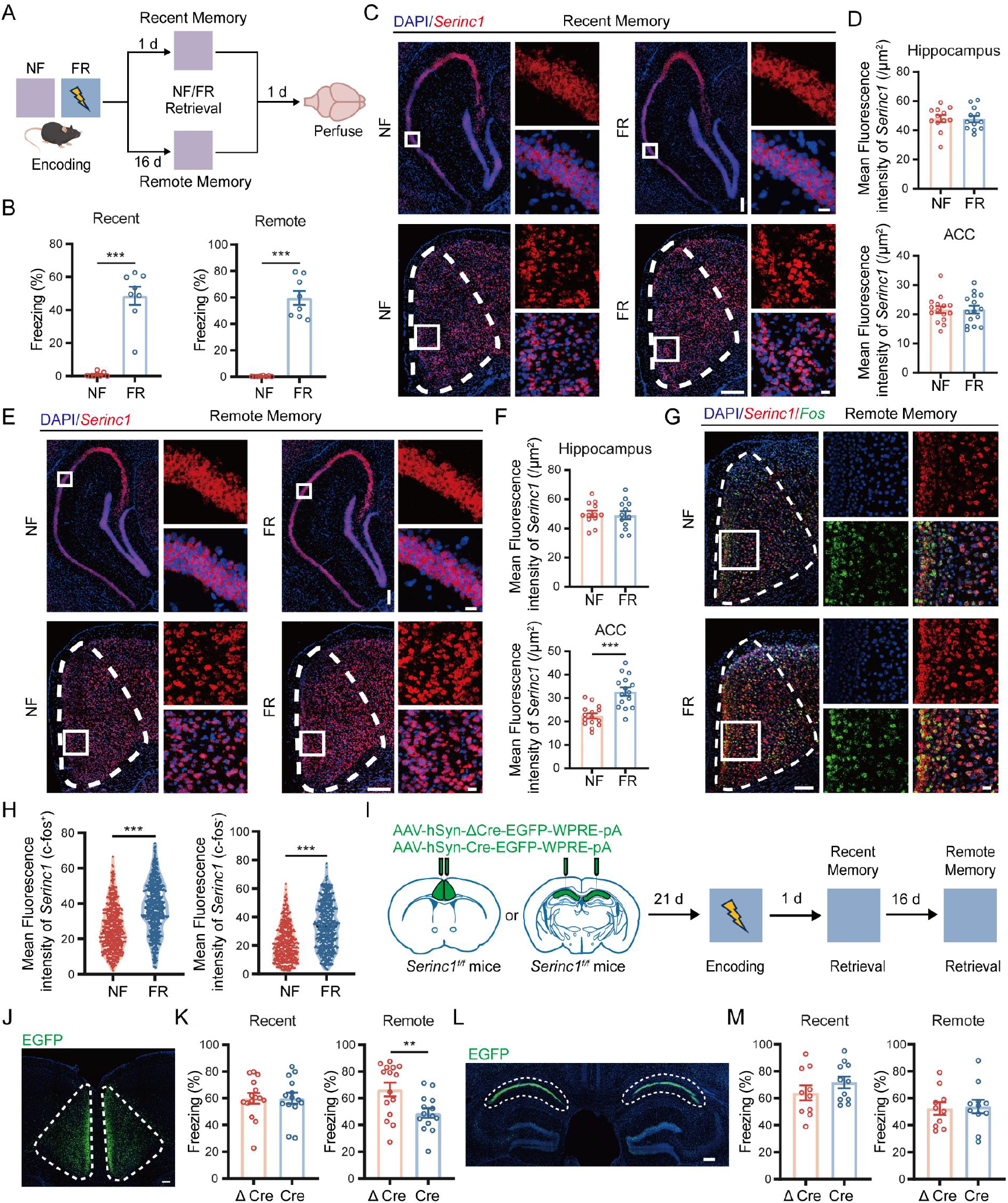
*Serinc1* is selectively induced in the ACC during remote memory and is required for remote memory retention. **(A)** Schematic of the experimental design for recent (Day 1) and remote (Day 16) memory retrieval following contextual fear conditioning (CFC), followed by RNAscope *in situ* hybridization. **(B)** Freezing behavior during retrieval testing 1 or 16 days post-training in fear-conditioned (FR) and unconditioned no-fear control (NF) mice. **(C** and **D)** Representative RNAscope images (C) and quantification (D) of *Serinc1* mRNA expression in the hippocampal CA1 and layer II/III of the anterior cingulate cortex (ACC) following recent (Day 1) memory retrieval. Scale bars, 200 μm (insets, 20 μm). **(E** and **F)** Representative RNAscope images (E) and quantification (F) of *Serinc1* mRNA expression in the hippocampal CA1 and layer II/III of the ACC following remote (Day 16) memory retrieval, showing selective induction in the ACC. Scale bars, 200 μm (insets, 20 μm). **(G)** Representative RNAscope images showing dual-fluorescence labeling of *Serinc1* mRNA (red) and *Fos* mRNA (green) in layer II/III of the ACC following remote memory retrieval. Scale bars, 100 μm (insets, 20 μm). **(H)** Single-cell quantification of *Serinc1* transcript abundance in active engram-associated (*Fos*⁺) cells and neighboring bystander (*Fos*⁻) cells within layer II/III of the ACC after remote memory retrieval. **(I)** Experimental design for region-specific conditional knockout of *Serinc1*. Adult *Serinc1^f/f^* mice received bilateral microinjections of AAV-Cre or control AAV-ΔCre into the ACC or hippocampal CA1, followed by CFC and memory testing at 1 or 16 days post-training. **(J)** Representative fluorescence image confirming bilateral viral targeting of the ACC in *Serinc1^f/f^* mice. Scale bar, 500 μm. **(K)** Freezing behavior during recent (Day 1) and remote (Day 16) memory retrieval following ACC-targeted deletion of *Serinc1*. **(L)** Representative fluorescence image confirming bilateral viral targeting of hippocampal CA1 in *Serinc1^f/f^* mice. Scale bar, 500 μm. **(M)** Freezing behavior during recent and remote memory retrieval following CA1-targeted deletion of *Serinc1*.

To resolve the cellular pattern of this neocortical induction, we harvested ACC tissue 15 min after remote memory retrieval and simultaneously quantified *Serinc1* and the immediate-early gene *Fos* (fig. S1, A and B). Multiplexed cellular profiling revealed that *Serinc1* was broadly elevated across both *Fos*^+^ engram neurons and *Fos^-^* bystander neurons within the ACC (Fig. 1, G and H). Parallel tracking in hippocampal CA1 confirmed a complete lack of induction in both populations (fig. S1, C and D), indicating that *Serinc1* operates as a network-wide neocortical modifier rather than an ensemble-restricted marker.

To establish the functional necessity of this cortex-restricted response, we generated *Serinc1* floxed mice (*Serinc1^f/f^*) and selectively deleted *Serinc1* in the adult ACC or hippocampal CA1 neurons via bilateral delivery of AAV-hSyn-Cre-EGFP (Fig. 1I). Adult-onset deletion of *Serinc1* in the ACC neurons produced no detectable impairment in recent fear memory (Day 1), demonstrating that local SERINC1 is dispensable for initial context encoding, shock responsiveness, and immediate retrieval (Fig. 1K). However, when re-tested at Day 16, ACC neuron-specific *Serinc1* knockout mice exhibited a severe and selective disruption of remote fear memory expression (Fig. 1K). Control assays confirmed that this remote memory deficit occurred independently of gross locomotor impairments or baseline anxiety in the elevated plus maze (fig. S2, A to G). Although ACC neuron-specific *Serinc1* knockout mice exhibited a modest increase thigmotaxis in the open-field, overall distance traveled and primary affective behavior remained unperturbed. Conversely, targeted deletion of *Serinc1* in the dorsal hippocampal CA1 neurons altered neither recent nor remote fear memory (Fig. 1, L and M). Together, these findings establish a double dissociation: while hippocampal neuronal SERINC1 is dispensable for recent and remote memory retention, ACC neuron-specific *Serinc1* expression serves as an essential molecular gatekeeper for systems memory consolidation.

### Neuronal SERINC1 maintains ACC synaptic density and transmission

To investigate the cellular mechanisms underlying this remote memory failure, we crossed *Serinc1^f/f^* mice with CMV-Cre transgenic mice to generate constitutive *Serinc1* knockout (*Serinc1^ko^*) animals (fig. S3, A to C). Global *Serinc1* deletion caused no abnormalities in body weight or brain morphology (fig. S3, D and E). Sparse EGFP labeling of primary cortical neurons (DIV14) derived from *Serinc1^f/f^* and *Serinc1^ko^* embryos revealed identical soma sizes, dendritic arborization, and axonal length (fig. S4, A to E), indicating that SERINC1 is not required for fundamental structural maturation.

However, co-immunostaining mature primary cortical networks for pre- and postsynaptic marker pairs—excitatory vGLUT1/Homer1 and inhibitory vGAT/Gephyrin—revealed an approximately 40% reduction in structural synapse density in *Serinc1^ko^* neurons (Fig. 2, A to D). Individual synaptic puncta dimensions were unaffected (fig. S5, A and B), indicating a selective reduction in total synapse number rather than synaptic scaling. Neuronal re-expression of SERINC1 via DIV3 AAV-hSyn-*Serinc1*-P2A-EGFP infection fully rescued excitatory and inhibitory synapse densities to wild-type levels (Fig. 2, E and F, and fig. S5, C to F). To further verify a cell-autonomous mechanism, neuron-specific deletion of *Serinc1* in cultured cortical neurons using AAV-hSyn-Cre-EGFP faithfully phenocopied the global knockout (fig. S6, A to F), solidifying the intrinsic requirement for neuronal SERINC1 in preserving synaptic structural integrity.

**Fig. 2.**
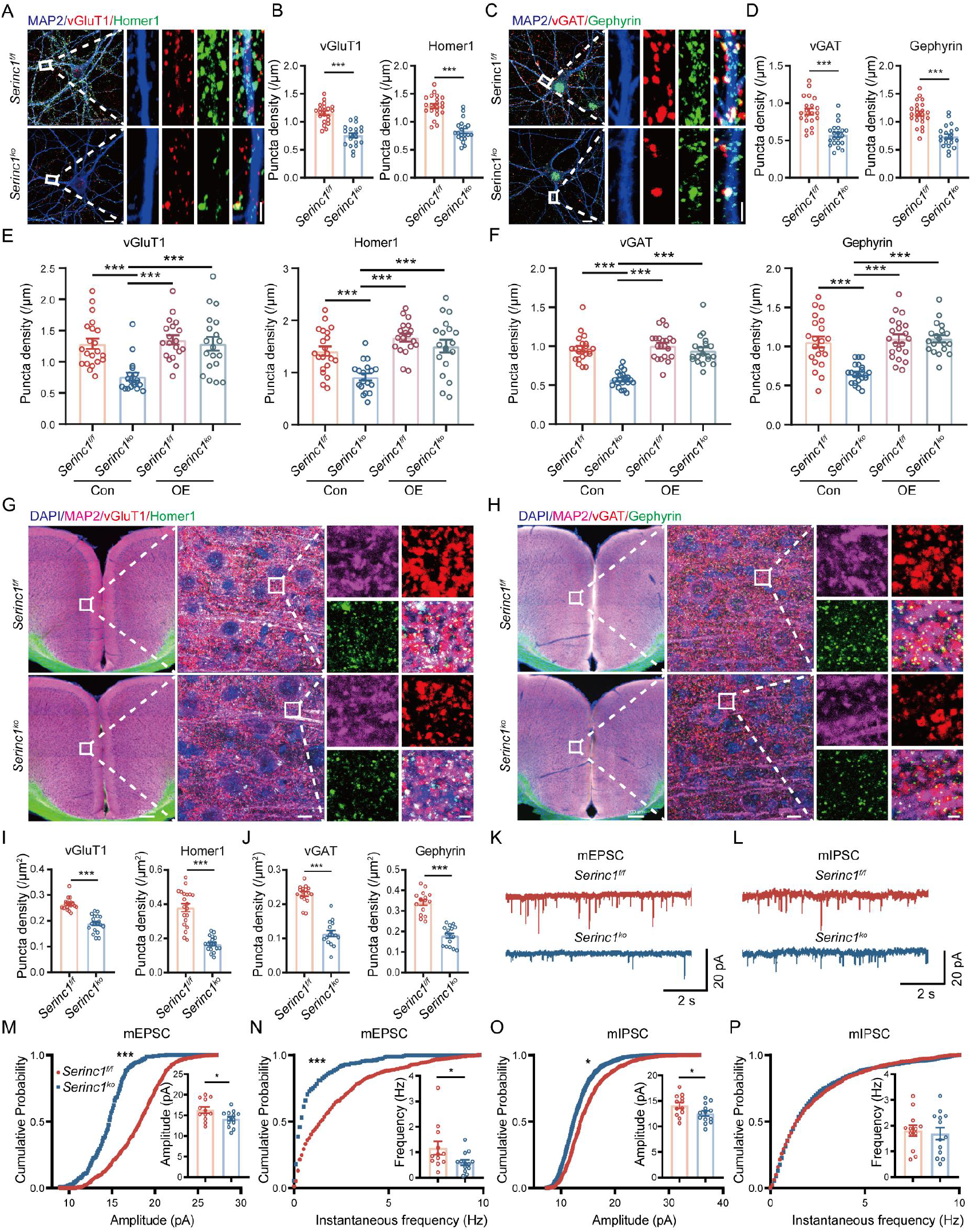
Neuronal SERINC1 maintains ACC synaptic density and electrophysiological function. **(A** and **B)** Representative confocal images (A) and quantification (B) of excitatory synaptic puncta labeled with presynaptic vGLUT1 and postsynaptic Homer1 in primary cortical neurons derived from *Serinc1^f/f^* and *Serinc1^ko^* mice. Scale bars, 10 μm (insets, 2 μm). **(C** and **D)** Representative confocal images (C) and quantification (D) of inhibitory synaptic puncta labeled with presynaptic vGAT and postsynaptic Gephyrin in primary cortical neurons derived from *Serinc1^f/f^* and *Serinc1^ko^* mice. Scale bars, 10 μm (insets, 2 μm). **(E** and **F)** Quantification of excitatory (E, vGLUT1/Homer1) and inhibitory (F, vGAT/Gephyrin) synaptic puncta densities in primary cortical neurons infected at DIV3 with control AAV-EGFP or rescue AAV-*Serinc1*-P2A-EGFP and immunostained at DIV14. **(G** to **J)** Representative images of excitatory (G) and inhibitory (H) synaptic puncta in layer II/III of the ACC of adult *Serinc1^f/f^* and *Serinc1^ko^* mice, with corresponding quantification of excitatory (I) and inhibitory (J) synaptic densities. Scale bars, 10 μm (insets, 2 μm). **(K** and **L)** Representative whole-cell patch-clamp recording traces of miniature excitatory postsynaptic currents (mEPSCs; K) and miniature inhibitory postsynaptic currents (mIPSCs; L) recorded from layer II/III pyramidal neurons in acute ACC slices from *Serinc1^f/f^* and *Serinc1^ko^* mice. **(M** to **P)** Cumulative probability distributions and summary plots of mEPSC amplitude (M), mEPSC frequency (N), mIPSC amplitude (O), and mIPSC frequency (P).

Histological and ultrastructural evaluations in adult *Serinc1^ko^* mice confirmed a concomitant reduction in excitatory (vGLUT1/Homer1) and inhibitory (vGAT/Gephyrin) synaptic puncta density in layer II/III of the ACC (Fig. 2, G to J, and fig. S5, G and H), alongside a lower physical synapse density assessed by transmission electron microscopy (fig. S7). Notably, active-zone length, synaptic vesicle density per bouton, and postsynaptic density thickness remained intact in remaining synapses (fig. S7). In line with our regional transcript mapping, hippocampal CA1 synaptic puncta density remained unaffected (fig. S8). Whole-cell patch-clamp recordings from ACC layer II/III pyramidal neurons in acute slices and cultured cortical networks revealed parallel electrophysiological deficits: *Serinc1^ko^* neurons exhibited significant reductions in miniature excitatory postsynaptic current (mEPSC) frequency and amplitude, as well as reduced miniature inhibitory postsynaptic current (mIPSC) amplitude (Fig. 2, K to P and fig. S9). While mIPSC amplitude was markedly reduced, mIPSC frequency remained unaltered, indicating that SERINC1 loss destabilizes postsynaptic receptor organization without altering basal presynaptic GABAergic release probability. Thus, neuronal SERINC1 maintains baseline synaptic density and functional transmission specifically within neocortical circuits.

### SERINC1 gates learning-induced, *de novo* callosal synaptogenesis during memory consolidation

Systems memory consolidation involves the progressive integration of neocortical networks, including interhemispheric callosal connections between bilateral ACC hubs (*24–28*). To determine whether this process engages *de novo* interhemispheric synaptogenesis, and whether it requires SERINC1, we performed mammalian mGRASP (GFP Reconstitution Across Synaptic Partners) imaging (*29–32*) (Fig. 3, A and B). AAV-CAG-pre-mGRASP-mCerulean was injected into the right ACC and AAV-CAG-post-mGRASP-tdTomato into the left ACC of *Serinc1^f/f^* and *Serinc1^ko^* mice. Tissue was collected 16 days after fear training (FT) without retrieval testing to isolate offline structural consolidation.

**Fig. 3.**
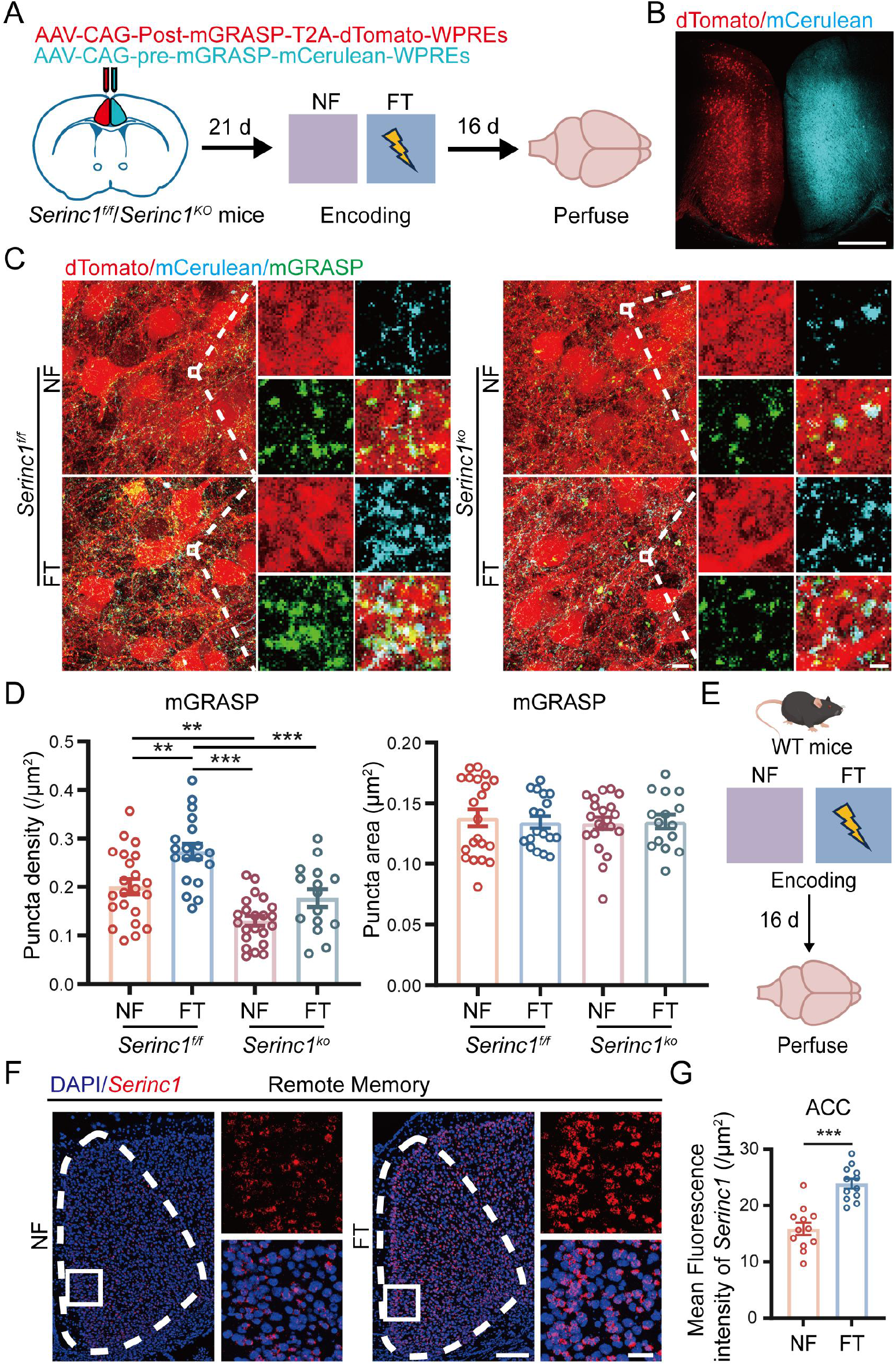
SERINC1 is required for learning-induced, *de novo* callosal synaptogenesis during memory consolidation. **(A)** Schematic of mammalian GFP reconstitution across synaptic partners (mGRASP) experimental strategy to map interhemispheric callosal connections. *Serinc1^f/f^* and *Serinc1^ko^* mice received unilateral microinjections of AAV-pre-mGRASP-mCerulean into the right ACC and AAV-post-mGRASP-tdTomato into the left ACC. After 21 days of viral expression, mice underwent CFC, and brains were harvested 16 days after fear training (FT) without retrieval testing to examine interhemispheric synapse assembly during memory consolidation. NF, no-fear controls. **(B)** Representative fluorescence coronal section showing targeted viral expression of pre-mGRASP-mCerulean (right ACC) and post-mGRASP-tdTomato (left ACC). Scale bar, 500 μm. **(C** and **D)** Representative confocal images (C) and quantification (D) of reconstituted interhemispheric mGRASP puncta (green) on postsynaptic dendrites of ACC neurons 16 days post-training. Quantification includes mGRASP puncta density and average puncta size. Scale bars, 10 μm (insets, 1 μm). **(E)** Schematic of the experimental design for measuring *Serinc1* transcript induction specifically during memory consolidation (harvested 16 days post-training without memory retrieval testing). **(F** and **G)** Representative RNAscope images (F) and quantification (G) of *Serinc1* mRNA expression in the ACC during memory consolidation. Scale bars, 200 μm (insets, 20 μm).

In fear-trained control *Serinc1^f/f^* mice, quantitative mGRASP imaging revealed a marked increase in reconstituted synaptic puncta along postsynaptic dendrites in the contralateral ACC compared with context-only controls (Fig. 3, C and D), providing direct evidence for learning-induced, *de novo* interhemispheric synaptogenesis during memory consolidation. Crucially, this experience-dependent callosal synaptogenesis was completely abolished in *Serinc1^ko^* mice (Fig. 3, C and D), demonstrating that SERINC1 is required for learning-induced, *de novo* circuit wiring. Furthermore, multiplexed RNAscope in fear-conditioned wild-type mice at Day 16 post-training revealed sustained elevation of *Serinc1* transcripts in layer II/III of the ACC in the absence of memory retrieval (Fig. 3, E to G), suggesting that associative learning triggers a persistent, offline upregulation of *Serinc1* expression that supports long-term structural circuit assembly.

### SERINC1 preserves PUFA-phospholipid pools and synaptic signaling during memory consolidation

To identify the biochemical mechanism linking SERINC1 to neocortical structural stability, we performed quantitative lipidomic profiling on micro-dissected ACC tissue harvested at Day 16 post-conditioning without retrieval (Fig. 4 and fig. S10). Comparing fear-conditioned wild-type mice (*Serinc1^f/f^*-FT) with unconditioned no-fear controls (*Serinc1^f/f^*-NF) demonstrated that physiological memory consolidation actively remodels the neocortical matrix toward a flexible, fluid landscape, enriching polyunsaturated fatty acid (PUFA)-bearing and ether-linked phospholipids—such as PE(O-18:1_22:3), PE(24:1_18:1), PC(44:3), and LPC(20:1)—while suppressing rigid sphingolipids, including ceramides and hexosylceramides (Fig. 4, A and B).

**Fig. 4.**
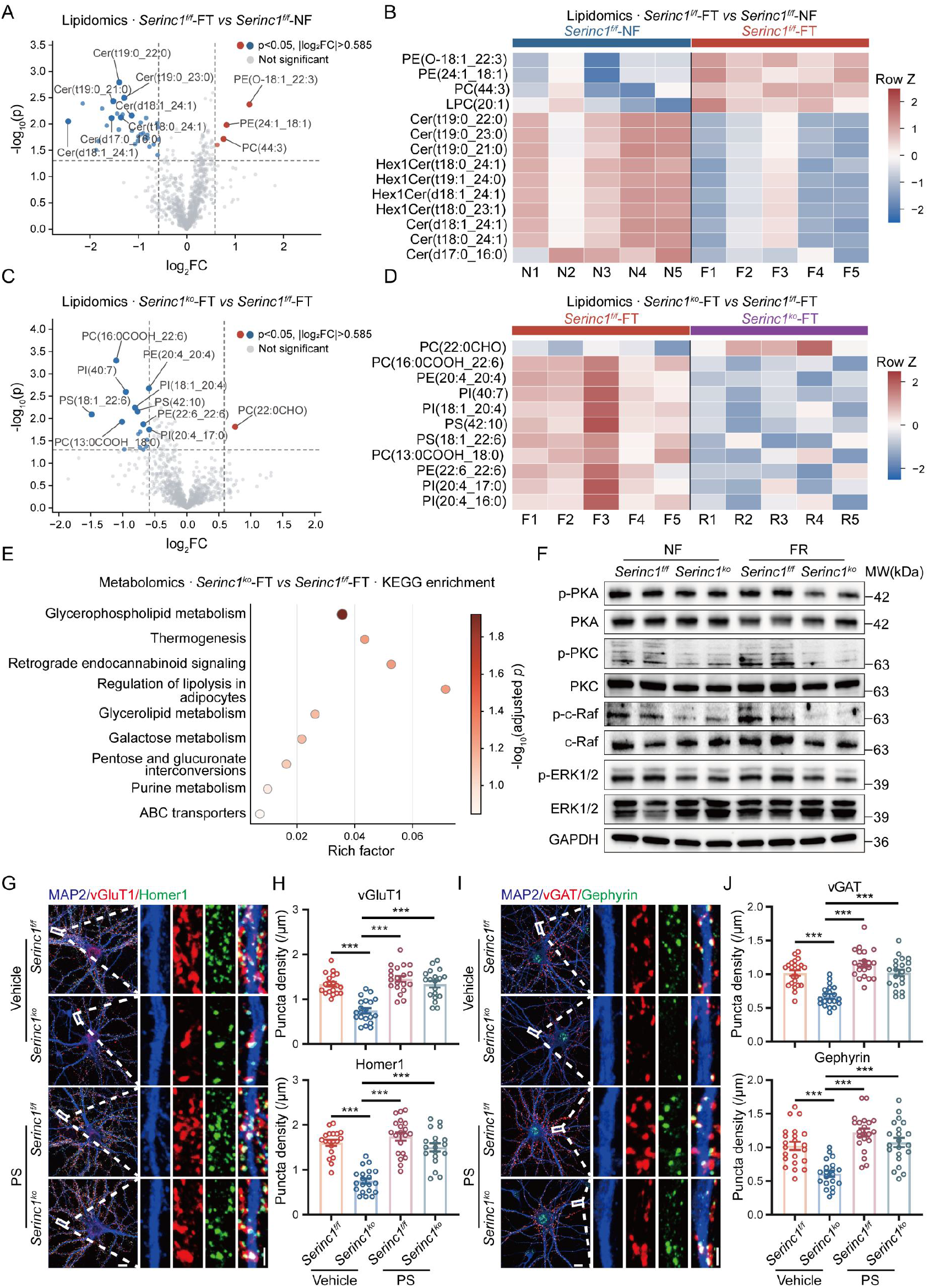
SERINC1 prevents activity-dependent collapse of anionic PUFA-phospholipid pools and preserves downstream synaptic signaling. **(A)** Volcano plot of positive-ion untargeted lipidomic profiling of ACC tissue from *Serinc1^f/f^* mice subjected to fear training (FT) versus unconditioned no-fear controls (NF) during memory consolidation. *P* < 0.05 (unadjusted two-sided Welch’s t-tests). FC, fold change; NS, not significant. Each point represents an individual lipid feature passing quality control (QC), blank, and missingness filters. **(B)** Heatmap of top differentially abundant lipid species identified in (A), comparing NF and FT *Serinc1^f/f^* cohorts. Abundance values were log_2_-transformed and displayed as row-wise Z-scores to highlight relative within-lipid pattern shifts. **(C)** Volcano plot of positive-ion lipidomic profiling of ACC tissue comparing FT-conditioned *Serinc1^f/f^* and *Serinc1^ko^* mice during memory consolidation (*P* < 0.05, unadjusted two-sided Welch’s t-tests). **(D**) Heatmap of differentially abundant phospholipid species during memory consolidation. Multiple phosphatidylserine (PS), phosphatidylcholine (PC), and phosphatidylinositol (PI) species were markedly depleted in *Serinc1^ko^* ACC tissue (log_2_-transformed abundance; row-wise Z-score). **(E)** KEGG pathway enrichment analysis of significantly altered metabolites between FT-conditioned *Serinc1^f/f^* and *Serinc1^ko^* ACC tissue. The x-axis indicates the rich factor, dot size represents the number of mapped compounds, and dot color indicates −log_10_ (adjusted *P* value). **(F)** Representative immunoblots of phosphorylated and total levels of PKA, PKC, c-Raf, and ERK1/2 (MAPK) in ACC lysates from *Serinc1^f/f^* and *Serinc1^ko^* mice under NF and fear recall (FR) conditions. GAPDH served as a loading control. **(G** and **H)** Representative confocal images (G) and quantification (H) of excitatory synaptic puncta density (vGLUT1/Homer1) in primary cortical neurons derived from *Serinc1^f/f^* and *Serinc1^ko^* mice following 48 h treatment (DIV12–DIV14) with vehicle or exogenous phosphatidylserine (PS, 20 μg/mL). Scale bars, 10 μm (insets, 2 μm). **(I** and **J)** Representative confocal images (I) and quantification (J) of inhibitory synaptic puncta density (vGAT/Gephyrin) in primary cortical neurons derived from *Serinc1^f/f^* and *Serinc1^ko^* mice following vehicle or exogenous PS treatment. Scale bars, 10 μm (insets, 2 μm).

While unconditioned *Serinc1^ko^* mice maintained baseline lipid profiles comparable to controls (fig. S10), fear-conditioned *Serinc1^ko^* mice (*Serinc1^ko^*-FT) exhibited a severe lipidomic collapse (Fig. 4, C and D). *Serinc1^ko^* ACC tissue failed to sustain PUFA-containing inner-leaflet signaling lipids, showing selective depletion of docosahexaenoic acid (DHA, 22:6)-bearing phosphatidylserines (PS(18:1_22:6) and PS(42:10)) and arachidonic acid (AA, 20:4)-bearing phosphatidylinositols (PI(18:1_20:4) and PI(20:4_17:0)), alongside an accumulation of oxidized species (PC(22:0 CHO)) (Fig. 4, C and D). Non-targeted polar metabolomics and KEGG enrichment confirmed major disruptions in glycerophospholipid metabolism and downstream retrograde endocannabinoid signaling (Fig. 4E and fig. S11).

Because inner-leaflet anionic lipids act as essential docking platforms for signaling and synaptic regulators (*17, 33–36*), we evaluated downstream synaptic kinase activation. ACC tissue from *Serinc1^ko^* mice collected after remote fear memory retrieval displayed significant reductions in phosphorylated PKC, MAPK, and PKA signaling (Fig. 4F).

Guided by our lipidomic identification of selective PS depletion, we tested whether restoring PS could causally reverse the synaptic deficits. In cultured primary *Serinc1^ko^* neurons, which exhibit baseline PS deficiency (fig. S12, A and B), exogenous supplementation with PS—but not sphingomyelin (SM)—fully rescued excitatory and inhibitory synaptic puncta densities (Fig. 4, G to J, and fig. S13). Thus, SERINC1 prevents activity-dependent exhaustion of anionic PUFA-phospholipid reserves, safeguarding downstream synaptic signaling and structural integrity.

### Neuronal restoration of SERINC1 in the ACC rescues synaptic integrity and remote memory in *Serinc1* knockout mice

Finally, we tested whether targeted re-expression of SERINC1 in ACC neurons is sufficient to restore synaptic structure and remote memory in global knockout mice. Adult *Serinc1^f/f^* and *Serinc1^ko^* mice received bilateral ACC injections of AAV-hSyn-*Serinc1*-P2A-EGFP or control AAV-hSyn-EGFP prior to CFC (Fig. 5, A and B).

**Fig. 5.**
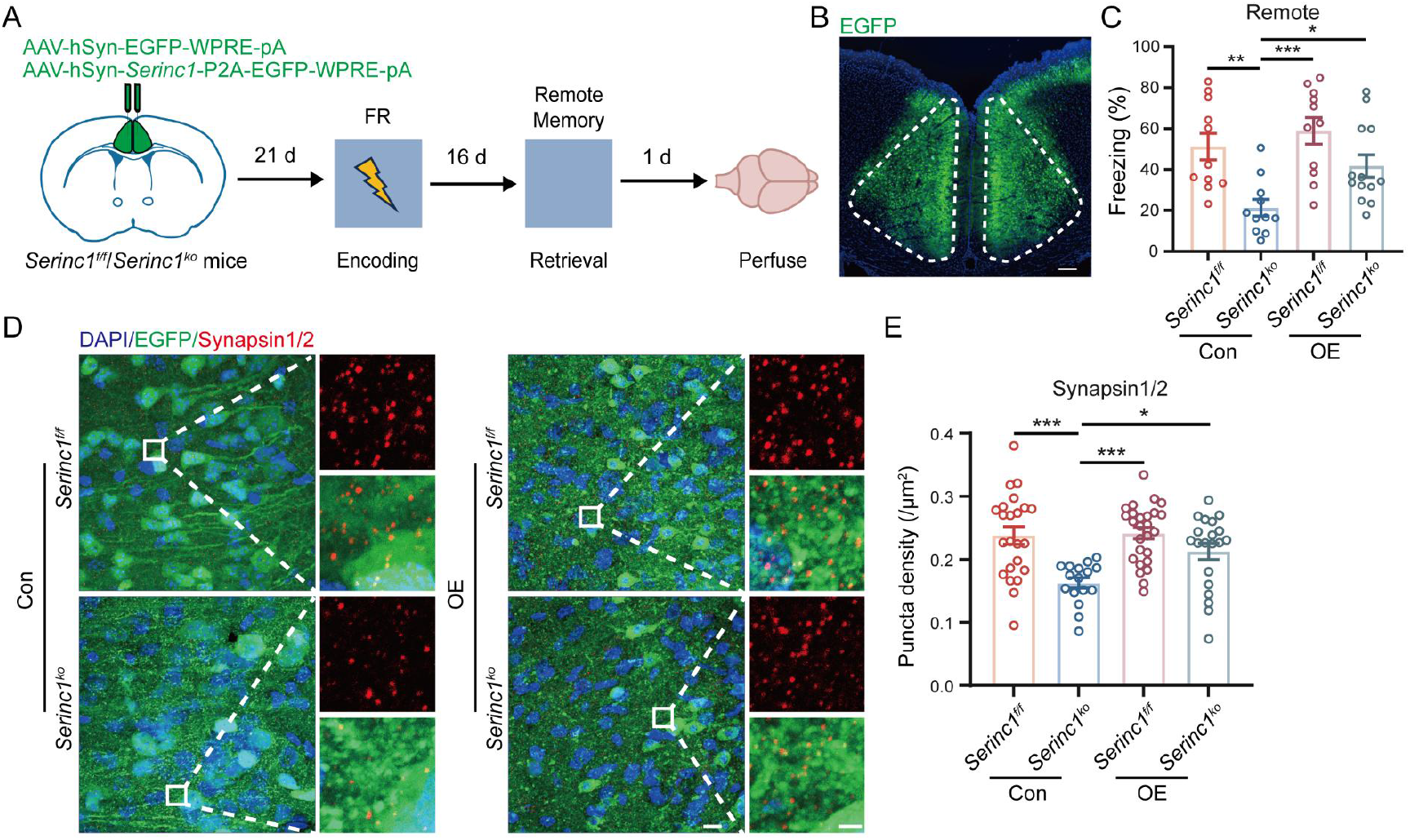
Neuronal SERINC1 restoration in the ACC rescues synaptic integrity and remote memory in *Serinc1* knockout mice. **(A)** Schematic of the experimental timeline for viral rescue in the ACC. Adult *Serinc1^f/f^* and *Serinc1^ko^* mice received bilateral ACC microinjections of control AAV-EGFP or rescue AAV-*Serinc1*-EGFP. Contextual fear conditioning was conducted 21 days post-injection, followed by remote memory retrieval testing 16 days post-training. **(B)** Representative fluorescence coronal image confirming bilateral viral targeting and reporter expression in the ACC. Scale bar, 500 μm. **(C)** Freezing behavior during remote memory retrieval testing in control- and SERINC1-restored *Serinc1^f/f^* and *Serinc1^ko^* cohorts. **(D** and **E)** Representative confocal immunofluorescence images (D) and quantification (E) of Synapsin1/2-positive presynaptic puncta density in layer II/III of the ACC across control and SERINC1-restored groups. Scale bars, 10 μm (insets, 2 μm).

Control-treated *Serinc1^ko^* mice displayed characteristic remote fear memory deficits at Day 16 post-conditioning (Fig. 5C). Targeted neuronal re-expression of SERINC1 in the ACC significantly increased freezing behavior in *Serinc1^ko^* mice, rescuing remote memory performance to levels comparable to *Serinc1^f/f^* controls (Fig. 5C). Immunofluorescence analysis demonstrated that SERINC1 re-expression simultaneously rescued Synapsin1/2-positive synaptic puncta density in ACC layer II/III (Fig. 5, D and E). These results demonstrate that SERINC1 expression specifically within ACC neurons is sufficient to maintain structural synapse density and gate neocortical memory consolidation.

## Discussion

Our findings establish that systems memory consolidation relies on a localized neocortical program that remodels the membrane lipid landscape to support long-term synaptic preservation. By deploying targeted genetic deletions and interhemispheric mGRASP imaging directly within the ACC, we demonstrate that the transmembrane regulator SERINC1 is required to sustain baseline synaptic densities and physically construct *de novo* callosal networks during the multi-week consolidation window. Tracking *Serinc1* expression uncovered a stark spatiotemporal divergence: it is robustly induced network-wide in the ACC during memory consolidation, but remains locked at baseline within the hippocampus. While hippocampal neuronal deletion yields no structural synaptic decay, the mature ACC uniquely relies on neuronal *Serinc1* for basal synaptic maintenance and recruits its activity-dependent induction as a dedicated metabolic switch required to scale its physical synaptic matrix and durably safeguard emerging memory traces.

Crucially, our multiplexed RNAscope mapping revealed that this upregulation occurs globally across both active *Fos^+^* engram and neighboring *Fos^-^* bystander cells within the ACC. This broad induction may reflect paracrine volume transmission or the cumulative footprint of millions of offline replay events occurring over the 16-day consolidation window. Importantly, this network-wide cortical remodeling does not imply a loss of behavioral specificity. Instead, the generalized enrichment of cellular PS establishes a permissive lipid field across the ACC, where structural specificity is dictated by active circuit inputs meeting this metabolically primed cellular landscape.

At the cellular level, this architectural stabilization shifts the framework of systems memory consolidation from a purely protein-driven event to one fundamentally gated by neural lipid dynamics. Unbiased lipidomic profiling revealed that physiological consolidation in wild-type mice restructures the ACC matrix toward a flexible state enriched in structural and polyunsaturated phospholipids, alongside ceramide suppression. In contrast, memory consolidation in *Serinc1^ko^* mice triggers a catastrophic exhaustion of inner-leaflet anionic signaling lipids—specifically docosahexaenoic acid (DHA)-bearing phosphatidylserines and arachidonic acid (AA)-bearing phosphatidylinositols—accompanied by accumulation of lipid oxidation products. This targeted phospholipid collapse is directly corroborated by untargeted polar metabolomics, where KEGG analysis identified glycerophospholipid metabolism and retrograde endocannabinoid signaling as the top disrupted pathways. General structural ether-lipids remain intact in knockout mice, demonstrating that SERINC1 acts as an indispensable homeostatic engine required to preserve inner-leaflet anionic pools against consolidation-induced exhaustion.

To explain why *Serinc1* knockout mice exhibit lipid depletion selectively during consolidation rather than at baseline, we propose a working mechanism of activity-dependent lipid buffering. Under basal conditions, low membrane turnover maintains relative parity between wild-type and *Serinc1*-deficient lipid profiles. During memory consolidation, however, we hypothesize that sustained neural firing and Ca^2+^ dynamics accelerate the turnover and consumption of inner-leaflet PUFA-containing PS and PI pools (*37–39*). In wild-type neurons, SERINC1 buffers against this localized consumption to maintain signaling lipid reserves; in its absence, uncoupled consumption leads to net lipid exhaustion. This metabolic failure directly accounts for the observed endocannabinoid pathway disruptions, as depletion of AA-bearing precursors—such as PI(18:1_20:4)—deprives neurons of the substrate required for dynamic endocannabinoid synthesis (*34, 40–42*), compounding the structural failure to construct and preserve remote memory networks.

By maintaining this polyunsaturated, anionic inner-leaflet matrix, SERINC1 sustains the localized surface charge and membrane curvature required to license downstream structural plasticity signaling cascades. Indeed, following remote memory recall, *Serinc1* deficiency leads to marked reductions in phosphorylated PKC, MAPK, and PKA pathways, establishing a direct link between membrane lipid preservation and kinase activation driving spine morphogenesis. Crucially, because exogenous PS supplementation directly restores synaptic marker density in cultured *Serinc1^ko^* neurons, and neuron-specific viral restoration in the ACC rescues local synapse density and remote behavioral retention in vivo, SERINC1-dependent phospholipid preservation operates as an obligate metabolic gate for neocortical synaptogenesis. Our data demonstrate that callosal synaptic endpoints depend on the dense inner-leaflet anionic matrix provided by SERINC1 to build and sustain functional interhemispheric connectivity.

The discovery that SERINC1 drives macro-circuit allocation reframes systems memory consolidation as a lipid-gated structural process. While the SERINC family is best known in virology for its antiviral restriction activity, with SERINC3 and SERINC5 perturbing viral membrane lipid asymmetry and impairing membrane fusion (*16, 43, 44*), our findings establish a parallel biophysical principle in the mammalian brain: neuronal SERINC1 actively remodels the postsynaptic lipid landscape to anchor structural signaling complexes necessary for *de novo* synaptogenesis. Notably, both cortical lipid homeostasis and distributed cortical circuits supporting remote memory are progressively disrupted in Alzheimer’s disease (*45–49*). Together with our findings that neuronal SERINC1 links membrane lipid remodeling to synaptic structural organization, these observations raise the possibility that failure of lipid-dependent membrane maintenance represents an underappreciated mechanism contributing to the destabilization of long-range memory networks. More broadly, the metabolic breakdown of this SERINC1-dependent structural program may highlight a critical vulnerability underlying memory loss in dementia. Ultimately, our study establishes that associative learning initiates a cortex-specific, network-wide neuro-lipidomic program that connects whole-cell metabolic scaling to the macro-scale construction of interhemispheric circuits.

## Acknowledgments

We thank the Bio-imaging Core Facilities of SZBL and SMART for imaging support and the Multi-omics Mass Spectrometry Core Facility of the Bio-Tech Center at SMART for LC-MS analysis. We also thank Professor Nobuhiko Yamamoto for discussions and critical reading of the manuscript.

## Author Contributions

X. J. conceived the project. X. J. and S. W. designed the experiments. S. W. performed mouse behavioral tests, RNAscope, primary neuron culture, Immunocytochemistry and immunohistochemistry. X. X. L. and H. C. performed stereotaxic surgeries. H. C. performed electrophysiological recordings and analysis. X. L. and S. W. conducted lipidomic and metabolomic experiments and analyses. S. W., H. C., X. L. performed statistical analyses. X. J. and S. W. wrote the manuscript with input from all authors. X. J. supervised the project.

## Data and materials availability

All data are available in the main text or the supplementary materials. Additional information and materials used in this study are available from the corresponding author upon reasonable request.

## Materials and Methods

### Mouse Breeding, Genotyping and Husbandry

Wild-type C57BL/6J mice, *Serinc1* cKO mice, and *Serinc1* KO mice were used in this study. Mice were weaned at postnatal day 21 (P21) and group-housed (fewer than 5 mice per cage) under a 12-h light/12-h dark cycle with ad libitum access to food and water at the Shenzhen Bay Laboratory Animal Center or the Shenzhen Medical Academy of Research and Translation Animal Facility. Unless otherwise indicated, all experiments were performed using male mice. Genotyping was performed in-house for all animals. Primary neuronal cultures were prepared from embryonic day 16–18 mouse. Brain tissue for histological and biochemical analyses was collected from mice at P42–49. All animal procedures were conducted in accordance with the Guide for the Care and Use of Laboratory Animals and were approved by the Institutional Animal Care and Use Committees (IACUCs) of Shenzhen Bay Laboratory and the Shenzhen Medical Academy of Research and Translation.

### Plasmids and virus generation

For all neuron-specific *Serinc1* knockout experiments, adeno-associated viruses (AAVs) expressing EGFP-Cre or EGFP-ΔCre under the control of the human synapsin (hSyn) promoter were obtained from Taitool Bioscience. For *Serinc1* overexpression experiments, AAVs expressing *Serinc1*-P2A-EGFP under the hSyn promoter were generated by BrainVTA. Briefly, recombinant AAVs were produced by triple-plasmid transfection of HEK293 cells using a transfer plasmid containing the transgene flanked by AAV2 inverted terminal repeats (ITRs), together with Rep/Cap and adenoviral helper plasmids. Viral particles were purified by cesium chloride density-gradient ultracentrifugation, and viral titers were determined by quantitative PCR (qPCR). For all mammalian GFP reconstitution across synaptic partners (mGRASP) experiments, AAVs expressing pre-mGRASP-mCerulean or post-mGRASP-tdTomato under the control of the CAG promoter were obtained from BrainVTA. Unless otherwise indicated, all AAVs were used at a final titer of 1 × 10¹² viral genomes (vg) ml⁻¹.

### Stereotaxic surgery

Stereotaxic viral injections were performed in mice at P28-30. Mice were anesthetized with isoflurane (2.5% for induction and 1.5% for maintenance) and secured in a stereotaxic frame. After the skull was exposed, bregma and lambda were aligned in the same horizontal plane (within ±0.03 mm). Small burr holes were drilled above the target coordinates, and viral vectors were delivered through a glass micropipette using a microsyringe pump. Following each injection, the micropipette was left in place for 10 min before being slowly withdrawn to minimize reflux.

For neuron-specific *Serinc1* deletion, 350 nL of AAV-hSyn-Cre-EGFP was bilaterally injected into either the anterior cingulate cortex (ACC; AP, +0.70 mm; ML, ±0.30 mm; DV, −1.50 mm) or the hippocampus (AP, −2.00 mm; ML, ±1.50 mm; DV, −1.40 mm) at a rate of 100 nL min⁻¹. Control mice received bilateral injections of the same volume of AAV-hSyn-ΔCre-EGFP.

For *Serinc1* overexpression, 350 nL of AAV-hSyn-*Serinc1*-P2A-EGFP was bilaterally injected into the ACC at 100 nL min⁻¹. Control mice received an equal volume of AAV-hSyn-EGFP.

For hippocampal mGRASP experiments, 60 nL of AAV-CAG-pre-mGRASP-mCerulean was injected into the left dorsal CA3 (dCA3; AP, −2.15 mm; ML, +2.35 mm; DV, −2.15 mm), and 60 nL of AAV-CAG-post-mGRASP-tdTomato was injected into the right dorsal CA1 (dCA1; AP, −2.00 mm; ML, −1.50 mm; DV, −1.25 mm) at 30 nL min⁻¹.

For ACC mGRASP experiments, 400 nL of AAV-CAG-pre-mGRASP-mCerulean was injected into the right ACC (AP, +0.70 mm; ML, +0.30 mm; DV, −1.50 mm), and 400 nL of AAV-CAG-post-mGRASP-tdTomato was injected into the left ACC (AP, +0.70 mm; ML, −0.30 mm; DV, −1.50 mm) at 100 nL min⁻¹.

Unless otherwise indicated, all subsequent behavioral, histological, and biochemical experiments were performed 3 weeks after viral injection. Viral expression and injection sites were verified by fluorescence microscopy after completion of the experiments, and only mice with correct viral targeting were included in the final analyses.

### Contextual fear conditioning

Contextual fear conditioning (CFC) was performed using a standard fear conditioning chamber. Mice were allowed to freely explore the chamber for 3 min before receiving three footshocks (0.5 mA, 2 s each) separated by 1-min intershock intervals. Following the final foot shock, mice remained in the chamber for an additional 1 min before being returned to their home cages. The chamber was thoroughly cleaned with 75% ethanol between animals to eliminate olfactory cues.

To assess recent and remote contextual fear memory, mice were re-exposed to the original conditioning chamber for 3 min at either 1 day or 16 days after training, respectively, without presentation of any footshocks or other external stimuli. Freezing behavior, defined as the complete absence of movement except for respiration, was recorded throughout the 3-min retrieval session and quantified as the percentage of total time spent freezing.

### Open field test

Locomotor activity and anxiety-like behavior were assessed using the open field test. The apparatus consisted of a square acrylic arena (40 × 40 × 40 cm), with a central zone (20 × 20 cm) defined for analysis. Mice were gently placed in one corner of the arena and allowed to freely explore for 10 min while being recorded by an overhead camera connected to an automated video-tracking system. The arena was cleaned with 75% ethanol between trials to eliminate olfactory cues. Total distance traveled was quantified as an index of spontaneous locomotor activity, whereas the time spent in the center zone was used as an index of anxiety-like behavior.

### Elevated plus maze

Anxiety-like behavior was further evaluated using the elevated plus maze (EPM). The maze consisted of two opposing open arms (30 × 5 cm) and two opposing closed arms (30 × 5 cm, enclosed by 15-cm-high walls) connected by a central platform (5 × 5 cm) and elevated 50 cm above the floor. At the beginning of each trial, mice were placed in the center of the maze facing the same open arm and allowed to freely explore for 10 min. The maze was cleaned with 75% ethanol between animals. The number of entries into the open arms and the time spent in the open arms were quantified.

### Y-maze test

Spatial working memory was assessed using the Y-maze spontaneous alternation task. The Y-maze consisted of three identical arms (35 × 5 × 15 cm) positioned 120° apart and connected by a central triangular area. At the beginning of each trial, mice were placed at the end of one arm facing away from the center and allowed to freely explore the maze for 10 min. The maze was cleaned with 75% ethanol between trials to eliminate olfactory cues. The sequence of arm entries was recorded, and spontaneous alternation performance was calculated as the percentage of successive entries into all three arms without repetition.

### RNA in situ hybridization

Target-specific probe pairs against *Serinc1* (564421-B1) and *Fos* (142811-B2) mRNAs were designed using a proprietary algorithm (Chinese Patent No. ZL202110581853.9) and synthesized by GD Pinpoease Biotech. Fresh brain tissues were embedded in optimal cutting temperature (OCT) compound, rapidly frozen, and stored at −80 °C until processing. Coronal cryosections (15 μm) were prepared using a cryostat (Leica CM1950). Multiplex fluorescent RNA in situ hybridization was performed using the PinpoRNA Multiplex Fluorescent RNA In Situ Hybridization Kit (Pinpoease Biotech, Cat# PIF1000/PIF2000) according to the manufacturer’s instructions. Briefly, sections were post-fixed in 4% PFA for 30 min at 4 °C, dehydrated sequentially in 50%, 70%, and 100% ethanol for 5 min each, and incubated in Pretreatment Solution A for 10 min at room temperature. Sections were then treated with Protease II at 40 °C for 15 min before hybridization with target-specific probes at 40 °C for 2 h. Hybridization signals were sequentially amplified through three rounds of signal amplification according to the manufacturer’s protocol, followed by horseradish peroxidase (HRP)-mediated tyramide signal amplification (TSA) using spectrally distinct fluorophores to visualize individual RNA targets. Nuclei were counterstained with DAPI where indicated. Fluorescence images were acquired using a Leica SP8 confocal microscope equipped with a fluorescence lifetime imaging microscopy (FLIM) module.

### Primary neuronal culture

Primary cortical and hippocampal neurons were prepared from embryonic day (E) 16–18 mouse embryos as previously described. Briefly, pregnant mice were euthanized, and E16–E18 embryos were collected. Embryonic hippocampi or cerebral cortices were rapidly dissected in ice-cold calcium-, magnesium-, and phenol red-free D-Hanks’ balanced salt solution (Sigma-Aldrich, Cat# H2387). Tissues were digested with 0.25% trypsin (Gibco, Cat# 25200072) at 37 °C for 12 min and gently dissociated into single-cell suspensions by passing through a 70-μm cell strainer. Dissociated neurons were plated onto poly-D-lysine-coated glass coverslips in 24-well plates and initially cultured in DMEM/F12 medium (Gibco, Cat# 11330032) supplemented with 10% fetal bovine serum (Gibco, Cat# A5669701). After 4 h, the culture medium was replaced with maintenance medium consisting of Neurobasal-A medium (Gibco, Cat# 10888022) supplemented with 2% B27 (Gibco, Cat# 17504044) and 1% GlutaMAX (Gibco, Cat# 35050061). Neurons were maintained at 37 °C in a humidified incubator with 5% CO₂. For viral transduction experiments, AAVs were added to the cultures at DIV3. Unless otherwise indicated, all neuronal analyses were performed at DIV14-DIV16.

### Morphological analysis

Primary cortical neurons were isolated from *Serinc1^f/f^* and *Serinc1^ko^* mice, sparsely transduced with AAV-hSyn-EGFP at DIV3. At DIV14, cells were washed once with PBS and fixed with 4% paraformaldehyde for 30 min at room temperature. After three additional washes with PBS, coverslips were mounted onto microscope slides. Images were acquired using an LSM 990 confocal microscope. Axons were imaged with a 10× objective, whereas dendritic morphology and somatic area were examined with a 20× objective. Axonal and dendritic arbors were reconstructed and analyzed using the Simple Neurite Tracer (SNT) plugin in Fiji (*50, 51*).

### Immunocytochemistry

Immunocytochemistry was performed on primary neurons at DIV14–DIV16. Neurons were gently rinsed once with prewarmed phosphate-buffered saline (PBS) and fixed with 4% PFA for 20 min at room temperature. After three washes with PBS, cells were permeabilized with PBS containing 0.3% Triton X-100 for 15 min and subsequently blocked with 5% bovine serum albumin (BSA) in PBS for 30 min at room temperature. Cells were incubated overnight at 4 °C with primary antibodies diluted in blocking buffer, including guinea pig anti-MAP2 (Synaptic Systems, Cat# 188004), rabbit anti-VGLUT1 (Synaptic Systems, Cat# 135302), mouse anti-HOMER1 (Synaptic Systems, Cat# 160011), rabbit anti-vGAT (Synaptic Systems, Cat# 131002), and mouse anti-Gephyrin (Synaptic Systems, Cat# 147011). After three washes with PBS, cells were incubated for 2 h at room temperature with the appropriate fluorophore-conjugated secondary antibodies diluted in blocking buffer, including goat anti-guinea pig Alexa Fluor 594 (Invitrogen, Cat# A-11076), goat anti-rabbit Alexa Fluor Plus 647 (Invitrogen, Cat# A-32733), and goat anti-mouse Alexa Fluor 488 (Abcam, Cat# ab150113). Following three additional washes with PBS, coverslips were mounted with Fluoromount-G mounting medium (SouthernBiotech, Cat# 0100-01) and imaged using a Zeiss LSM 980 confocal microscope.

### Immunohistochemistry

Mice were deeply anesthetized with isoflurane and transcardially perfused with ice-cold PBS, followed by ice-cold 4% PFA in PBS. Brains were dissected, post-fixed overnight in 4% PFA at 4 °C, and washed three times with PBS. Free-floating horizontal sections (40 μm) were prepared using a vibratome (Leica VT1200S). Sections were blocked for 3 h at room temperature in blocking buffer consisting of PBS supplemented with 5% normal goat serum (NGS) and 0.4% Triton X-100. Sections were then incubated overnight at 4 °C with primary antibodies diluted in incubation buffer (PBS containing 5% NGS and 0.1% Triton X-100), including guinea pig anti-MAP2 (Synaptic Systems, Cat# 188004), rabbit anti-VGLUT1 (Synaptic Systems, Cat# 135302), mouse anti-HOMER1 (Synaptic Systems, Cat# 160011), rabbit anti-vGAT (Synaptic Systems, Cat# 131002), mouse anti-Gephyrin (Synaptic Systems, Cat# 147011), and chicken anti-Synapsin1/2 (Synaptic Systems, Cat# 106006). After three washes with PBS (10 min each), sections were incubated for 2 h at room temperature with the appropriate fluorophore-conjugated secondary antibodies diluted in PBS containing 5% NGS, including goat anti-guinea pig Alexa Fluor 594 (Invitrogen, Cat# A-11076), goat anti-rabbit Alexa Fluor Plus 647 (Invitrogen, Cat# A-32733), goat anti-mouse Alexa Fluor 488 (Abcam, Cat# ab150113), and goat anti-chicken Alexa Fluor Plus 647 (Invitrogen, Cat# A-32933). Sections were subsequently washed three times with PBS (10 min each), mounted onto glass slides, air-dried, and coverslipped with an antifade mounting medium containing DAPI (Solarbio, Cat# S2110). Fluorescence images were acquired using a Zeiss LSM 980 confocal microscope.

### Electrophysiology

Mice at 3-4 weeks of age were deeply anesthetized with tribromoethanol and quickly decapitated. Brain slices (300 µm) were made using a vibratome (Leica VT1200S) in an ice-cold cutting solution with the following composition: 2 mM KCl, 1.25 mM NaH_2_PO_4_, 7 mM MgSO_4_·7H_2_O, 0.5 mM CaCl_2_·2H_2_O, 26 mM NaHCO_3_, 10 mM Glucose, 250 mM Sucrose (pH 7.4, 300-310 mOsm). Brain slices were transferred to a chamber and incubated in artificial cerebrospinal fluid (ACSF; 124 mM NaCl, 3 mM KCl, 1.24 mM NaH₂PO₄, 2 mM MgSO₄·7H₂O, 11 mM glucose, 26 mM NaHCO₃, and 2 mM CaCl₂·2H₂O) for 30 min at 33°C, followed by incubation at room temperature for an additional 30 min before recordings were performed. All solutions were continuously bubbled with 5% CO₂/95% O₂.

For whole-cell recording, miniature excitatory postsynaptic currents (mEPSCs) and miniature inhibitory postsynaptic currents (mIPSCs) were recording at holding potentials of -70 mV and 0 mV, respectively. All recording were made form neurons in layer 2/3 of ACC and primary cortical neurons. The neurons were patched with 4-7 MΩ borosilicate glass pipettes. The perfusion (1.5-2 ml/min) using ACSF and supplemented with 1 µM tetrodotoxin (TTX) for mEPSC and mIPSC recording. The pipettes were filled with internal solution containing: 140 mM CsCH_3_SO_3_, 2 mM MgCl_2_·6H_2_O, 5 mM TEA-Cl, 10 mM HEPES, 1 mM EGTA, 2.5 mM Mg-ATP, 0.3 mM Na-GTP (pH 7.2-7.3 adjusted with CsOH, 290-300 mOsm). Synaptic current were sampled at 20 kHz and filtered at 3kHz. All recording were performed at room temperature. The cells with series resistance >30 MΩ were excluded from the analysis. Data were collected with a MultiClamp 700B amplifier and analyzed with Clampfit 11.2 software (Molecular Devices).

### Scanning electron microscopy (SEM)

ACC tissue from *Serinc1^f/f^* and *Serinc1^ko^* mice was analyzed by scanning electron microscopy (SEM). Samples were prepared as described (*52*). Further sample processing was performed according to standard operating procedures by the Bioimaging Core Facility at the Shenzhen Bay Laboratory. Images were acquired using a ZEISS GeminiSEM 360 scanning electron microscope.

### mGRASP

Mammalian GFP Reconstitution Across Synaptic Partners (mGRASP) was used to examine callosal synaptic connectivity between the bilateral ACCs during memory consolidation (*29–32*). AAV-CAG-pre-mGRASP-mCerulean, encoding the presynaptic mGRASP component, was unilaterally injected into the right ACC, whereas AAV-CAG-post-mGRASP-2A-tdTomato, encoding the complementary postsynaptic component, was unilaterally injected into the left ACC. After 21 d of viral expression, *Serinc1^f/f^* and *Serinc1^ko^* mice underwent fear training (FT) or a no-fear control procedure (NF). NF mice received matched handling and context exposure without footshock. To isolate offline memory consolidation from retrieval-induced effects, no retrieval test was performed, and tissue was collected directly 16 d after fear training. Mice were transcardially perfused with buffered saline followed by 4% paraformaldehyde, and brains were coronally sectioned at 40 μm using a vibratome. Images were acquired using a Leica FLIM 8 laser-scanning confocal microscope. Unless otherwise indicated, native mGRASP-GFP fluorescence was imaged without anti-GFP antibody amplification.

### Lipidomic profiling

ACC tissue samples were snap-frozen in liquid nitrogen and stored at -80°C. Approximately 5 mg of tissue was homogenized with 50 μl water at −50°C for 15 min, followed by 228 μl of ice-cold methanol containing the lipidomic and metabolomic internal-standard mixtures and 8 μl of BHT solution (1 mg/ml). MTBE (760 μl) was added and vortexed for 10 s; 230 μl water was then added and vortexed for 20 s. Samples were ultrasonicated for 10 min, incubated at −20°C for 30 min and centrifuged at 20,000 × g for 10 min at 6°C. The upper organic phase was collected for lipidomic analysis, whereas the lower phase was collected separately for polar metabolomic analysis. The upper organic phase was dried under a gentle stream of nitrogen or by vacuum centrifugation. Lipidomic extracts were sequentially reconstituted with 20 µL of dichloromethane/methanol (2:1, v/v) and then with 40 µL of acetonitrile/isopropanol/water (65:30:5, v/v/v) containing 5 mM ammonium acetate, each vortexed for 30 s, followed by addition of 10 µL of lipid internal standard, composed of 0.2 µg/ml of myristic acid-d27, LPC(16:0-d31), PG(16:0/18:1-d31), PE(16:0/18:1-d31), and C24:1-ceramide. The pool of 70 µL was centrifuged at 20,000 × g for 10 min at 6°C, and the supernatant was transferred to autosampler vials for LC–MS analysis. Lipid extracts were injected at 2 μl per run. Lipidomic analysis was performed on a Vanquish Flex UHPLC system coupled to an Orbitrap 480 mass spectrometer equipped with an OptaMax NG ion source (Thermo Fisher Scientific). Separation used a Waters ACQUITY UPLC BEH C18 column (100 Å, 1.8 μm, 2.1 × 100 mm). Mobile phase A was 60% acetonitrile in water and B was isopropanol/acetonitrile (9:1, v/v), both containing 10 mM ammonium acetate. The gradient was 20% B (0–0.5 min), 20–40% (0.5–1.5 min), 40–60% (1.5–3 min), 60–98% (3–13 min), 98% (13–16 min) and 20% (16.1–20 min). Full MS was acquired in both ion modes over m/z 100–1500 at 60,000 FWHM with standard AGC and 100-ms maximum injection time. DDA HCD MS/MS was acquired at 15,000 FWHM with stepped normalized collision energies of 20, 40 and 60%, standard AGC and 22-ms maximum injection time; spray voltages were +3.5 and −3.0 kV. Raw lipidomic files were processed using Compound Discoverer 3.5 for peak extraction, alignment, filtering and putative identification. Features were retained according to the platform-validated peak-quality and detection criteria. Lipid annotations were reported using LIPID MAPS shorthand. Structural claims were limited to the level supported by the available precursor and MS/MS evidence.

### Untargeted metabolomic profiling

The lower phase generated during the biphasic extraction was collected separately for polar metabolomic analysis, dried by vacuum centrifugation, and reconstituted in 30 μl of 25% acetonitrile. The metabolomic internal-standard mixture added during the initial extraction contained acetylcarnitine-d3 (0.3 μg/ml), decanoyl-L-carnitine-d3 (0.15 μg/ml), L-palmitoylcarnitine-d3 (0.15 μg/ml), L-leucine-d3 (1 μg/ml), L-phenylalanine-d5 (5 μg/ml), L-tryptophan-d5 (3.2 μg/ml), cholic acid-d4 (1.5 μg/ml), and chenodeoxycholic acid-d4 (1.5 μg/ml). Metabolomics was performed using a Nexera LC40DX3 UHPLC system coupled to a Zeno TOF 7600 mass spectrometer. Separation used a Waters ACQUITY UPLC HSS T3 column (100 Å, 1.8 μm, 2.1 × 100 mm) with water and acetonitrile, each containing 0.1% formic acid.

Positive-mode gradient: 5% B for 1 min, 5–40% B over 1 min, 40–100% B over 8 min, 100% B for 4 min, return to 5% B at 12.1 min and equilibrate for 2 min. Negative mode was identical except for 2% B at the start and return. Flow rate was 0.35 ml/min and autosampler temperature 4°C. IDA spectra were acquired over m/z 50–1500; ion-spray voltage was +5.5/−4.5 kV, source temperature 550°C, GS1/GS2/curtain gas/CAD 55/55/35/7 psi, and tandem-MS collision energy 35 eV with ±15-eV spread. Metabolite identification was performed using the HMDB, METLIN, and KEGG databases with a mass accuracy threshold of <5 ppm. Raw metabolomic files were processed using MExplorerUltimate for peak extraction, alignment, filtering and putative identification. Features missing in more than 50% of all samples were removed; remaining missing values were imputed as one-fifth of the minimum intensity for that metabolite. PCA was used for unsupervised visualization. PLS-DA/OPLS-DA models were evaluated by cross-validation and permutation testing, and model performance was reported using R² and Q².

### PS staining

Phosphatidylserine (PS) was detected using Annexin V-mCherry (Beyotime, Cat# C1069L). Primary neurons were gently rinsed once with prewarmed phosphate-buffered saline (PBS) and fixed with 4% paraformaldehyde for 20 min at room temperature. Following three washes with PBS, cells were incubated with 195 μl of Annexin V-mCherry Binding Buffer containing 5 μl of Annexin V-mCherry reagent for 1 h at room temperature in the dark. After staining, coverslips were mounted with DAPI Fluoromount-G mounting medium (SouthernBiotech, Cat# 0100-20). Fluorescence images were acquired using an Olympus FV3000 laser-scanning confocal microscope.

### Quantitative RT-PCR

To determine the efficiency of *Serinc1* deletion, total RNA was isolated from primary neuronal cultures infected with Cre- or ΔCre-expressing AAVs at DIV3-4 and harvested at DIV14 using the FastPure Cell/Tissue Total RNA Isolation Kit (Vazyme) according to the manufacturer’s instructions. RNA concentration and purity were determined using a NanoDrop ND-1000 spectrophotometer (Thermo Fisher Scientific). Complementary DNA (cDNA) was synthesized using the HiScript III All-in-one RT SuperMix Perfect for qPCR kit (Vazyme, R333-01). Quantitative real-time PCR (qRT-PCR) was performed using Taq Pro Universal SYBR qPCR Master Mix (Vazyme, Q712-02) on a 7900HT Fast Real-Time PCR System (Applied Biosystems) according to the manufacturer’s instructions. Relative *Serinc1* mRNA expression levels were normalized to Actb (β-actin) using the comparative Ct (2^−ΔΔCt) method. The primer sequences were as follows: *Serinc1*, forward 5′-TGTATCGCCTGTGCTTTGGT-3′ and reverse 5′-TGGGATGAAGAATGCGCCAA-3′. The Actb primer set was purchased from Sangon Biotech (B661302).

### Immunoblotting

ACC tissues were homogenized in RIPA lysis buffer (Solarbio, Cat# R0010) supplemented with protease and phosphatase inhibitor cocktail (Solarbio, Cat# P1261). Lysates were incubated on ice for 30 min and centrifuged at 15,000 × *g* for 10 min at 4 °C. Protein samples were separated by SDS–PAGE using precast gels (ACE Biotechnology, Cat# ET15412Gel) and transferred onto polyvinylidene difluoride (PVDF) membranes (Millipore, Cat# IPVH00010). Membranes were blocked for 30 min at room temperature using QuickBlock Blocking Buffer (Abmart, Cat# AWB-6017M) and incubated overnight at 4 °C with primary antibodies diluted in QuickBlock Western Primary Antibody Dilution Buffer (Beyotime, Cat# P0256). Primary antibodies included phospho-ERK1/2 (Thr202/Tyr204; Cell Signaling Technology, Cat# 5558), total ERK1/2 (Cell Signaling Technology, Cat# 137F5), phospho-PKC (pan, Ser660; Cell Signaling Technology, Cat# 9371), total PKC (Proteintech, Cat# 12919-1-AP), phospho-c-Raf (Ser338; Cell Signaling Technology, Cat# 9427), total c-Raf (Cell Signaling Technology, Cat# 9422), phospho-PKA Cα (Thr197; Cell Signaling Technology, Cat# 5661), and total PRKACA (Proteintech, Cat# 27398-1-AP). All primary antibodies were used at a dilution of 1:1,000. Following incubation with horseradish peroxidase (HRP)-conjugated secondary antibodies for 1 h at room temperature, immunoreactive bands were visualized using BeyoECL Star Chemiluminescent Substrate (Beyotime, Cat# P0018AM) and imaged with a ChemiDoc Imaging System (Bio-Rad).

### Confocal Image Acquisition and Analysis

Images of primary neuronal morphology were acquired using a Zeiss LSM 990 confocal microscope equipped with 10× or 20× objectives. Z-stack images were collected at 0.6 μm intervals, and maximum-intensity projections were generated from the six consecutive optical sections exhibiting the strongest fluorescence signals. For the analysis of synaptic puncta in primary neurons and the mouse ACC, images were acquired using Zeiss LSM 980 or LSM 990 confocal microscopes equipped with a 63× objective. Z-stack images were collected at intervals of 0.6 μm (primary neurons) or 1.5 μm (ACC), and maximum-intensity projections were generated from the six or ten consecutive optical sections with the strongest fluorescence signals, respectively. For PS staining in primary neurons, images were acquired using Olympus FV3000 confocal microscopes equipped with a 40× objective. Z-stack images were collected at intervals of 0.6 μm, and maximum-intensity projections were generated from the six consecutive optical sections with the strongest fluorescence signals, respectively. For mGRASP imaging, images were acquired using a Leica FLIM 8 confocal microscope equipped with 20× or 100× objectives. Z-stack images were collected at 1.5 μm intervals, and maximum-intensity projections were generated from the ten consecutive optical sections with the strongest fluorescence signals. Laser power, detector gain, and acquisition settings were optimized for each fluorescence channel and maintained constant across all biological replicates within each experiment. Identical staining procedures and imaging parameters were used for all samples within the same experiment. Background fluorescence was measured for each experimental group, and a uniform background threshold was applied during image processing. Quantitative image analyses were performed using ImageJ (NIH).

### Statistical analysis

Statistical analyses were performed using GraphPad Prism 10.0 (GraphPad Software). Comparisons between two groups were performed using unpaired two-tailed Student’s t-tests. Comparisons among multiple groups were performed using one-way or two-way analysis of variance (ANOVA). For lipidomic and metabolomic datasets, differential abundance was analyzed separately using two-sided Welch’s t-tests with Benjamini–Hochberg correction. Data are presented as mean ± SEM from at least three independent experiments. No statistical methods were used to predetermine sample size. Statistical significance was defined as *P* < 0.05. Unless otherwise indicated, significance levels are denoted as *P* < 0.05 (*), *P* < 0.01 (**), and *P* < 0.001 (***).

---

**Fig. S1.**
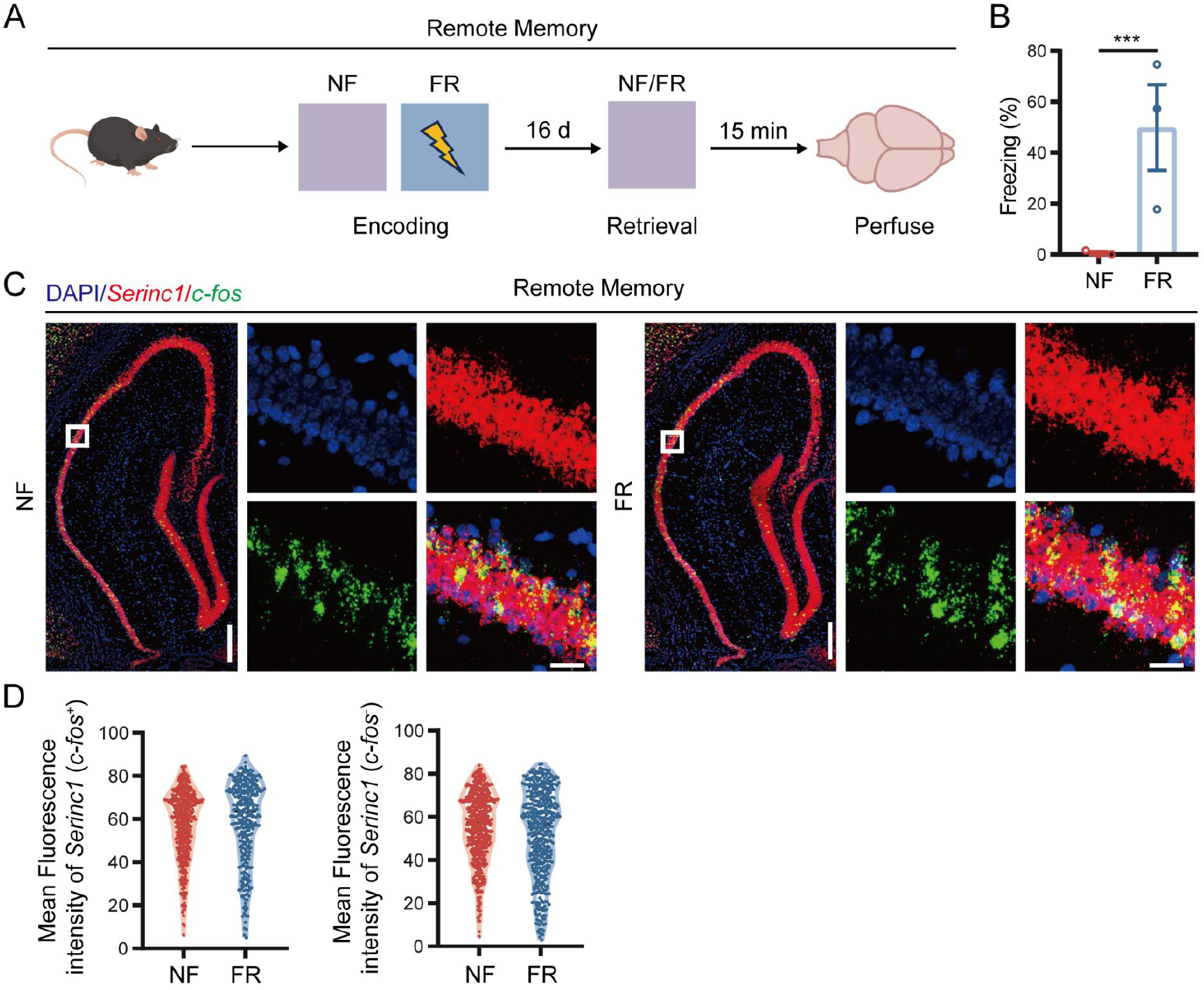
*Serinc1* expression is unchanged in hippocampal CA1 in remote memory. **(A)** Schematic of the experimental design. Mice underwent CFC to generate remote fear memory, and brains were collected 15 min after fear retrieval (FR) at Day 16 for RNAscope analysis of *Serinc1* and *Fos* mRNA expression. **(B)** Freezing behavior during retrieval testing in mice 16 days after fear conditioning. **(C)** Representative RNAscope images showing the colocalization of *Serinc1* mRNA (red) and *Fos* mRNA (green) in the hippocampus following remote memory retrieval. Zoomed in show the CA1 region. Scale bars, 200 μm (insets, 20 μm). **(D)** Single-cell quantification of *Serinc1* transcript abundance in active engram-associated (*Fos⁺*) neurons and neighboring bystander (*Fos⁻*) neurons in the hippocampal CA1 following remote memory retrieval.

**Fig. S2.**
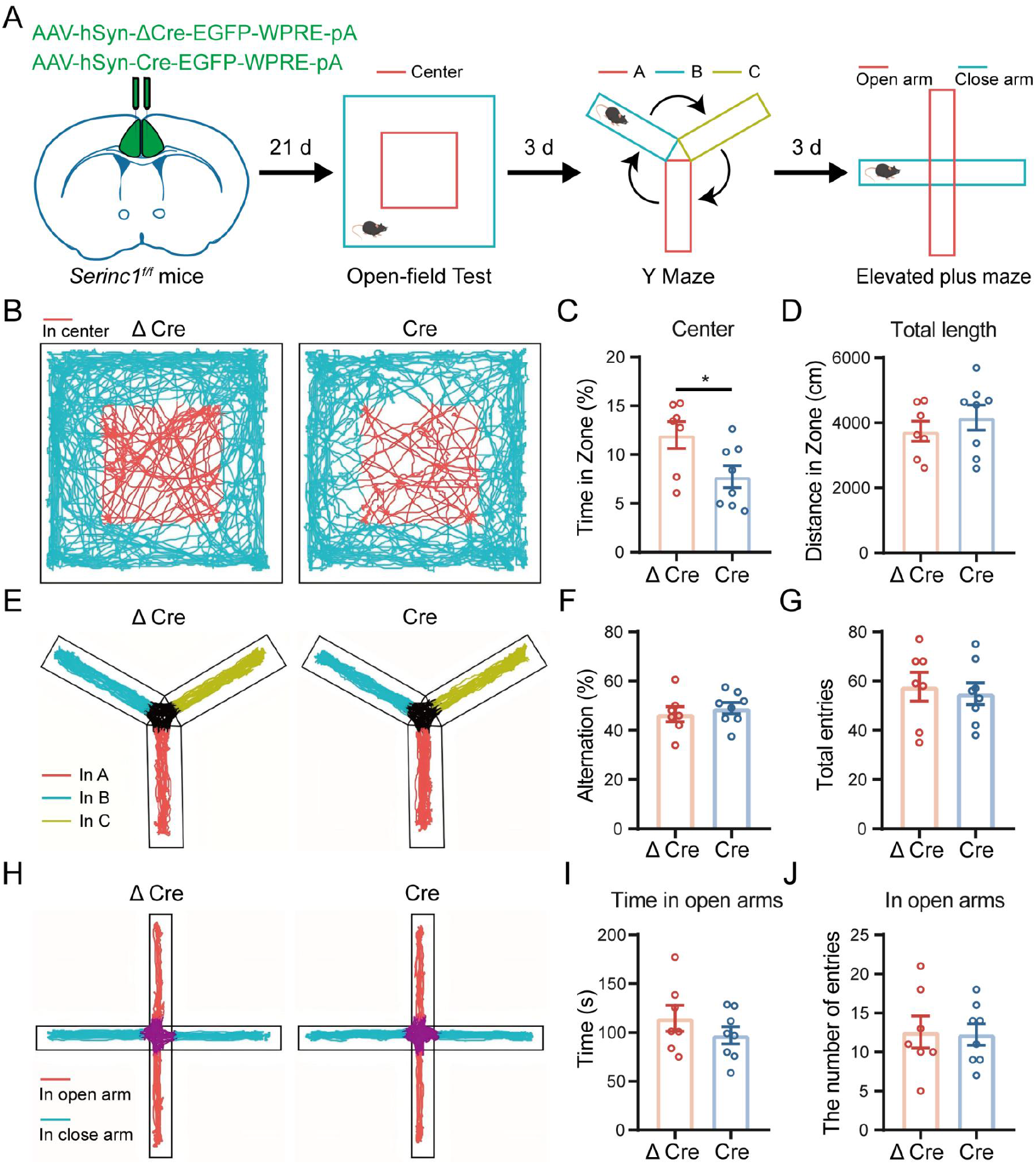
Behavioral characterization of mice with ACC neuron-specific deletion of *Serinc1*. **(A)** Schematic of the experimental design. Adult *Serinc1^f/f^* mice received bilateral stereotaxic injections of Cre- or ΔCre-expressing AAVs into the ACC to achieve ACC-specific *Serinc1* deletion. Mice subsequently underwent the open field test (OFT), Y-maze test, and elevated plus maze (EPM), with a 3-day interval between consecutive behavioral assays. **(B)** Representative locomotor trajectories in the OFT. Red and cyan indicate the center and peripheral zones, respectively. **(C** and **D)** Quantification of the percentage of time spent in the center zone (C) and the total distance traveled (D) in the OFT. **(E)** Representative exploration trajectories in the Y-maze test. Red, cyan, and yellow indicate arms A, B, and C, respectively. **(F** and **G)** Quantification of spontaneous alternation (F) and the total number of arm entries (G) in the Y-maze test. **(H)** Representative locomotor trajectories in the EPM. Red and cyan indicate the open and closed arms, respectively. **(I** and **J)** Quantification of the time spent in the open arms (I) and the number of open-arm entries (J) in the EPM.

**Fig. S3.**
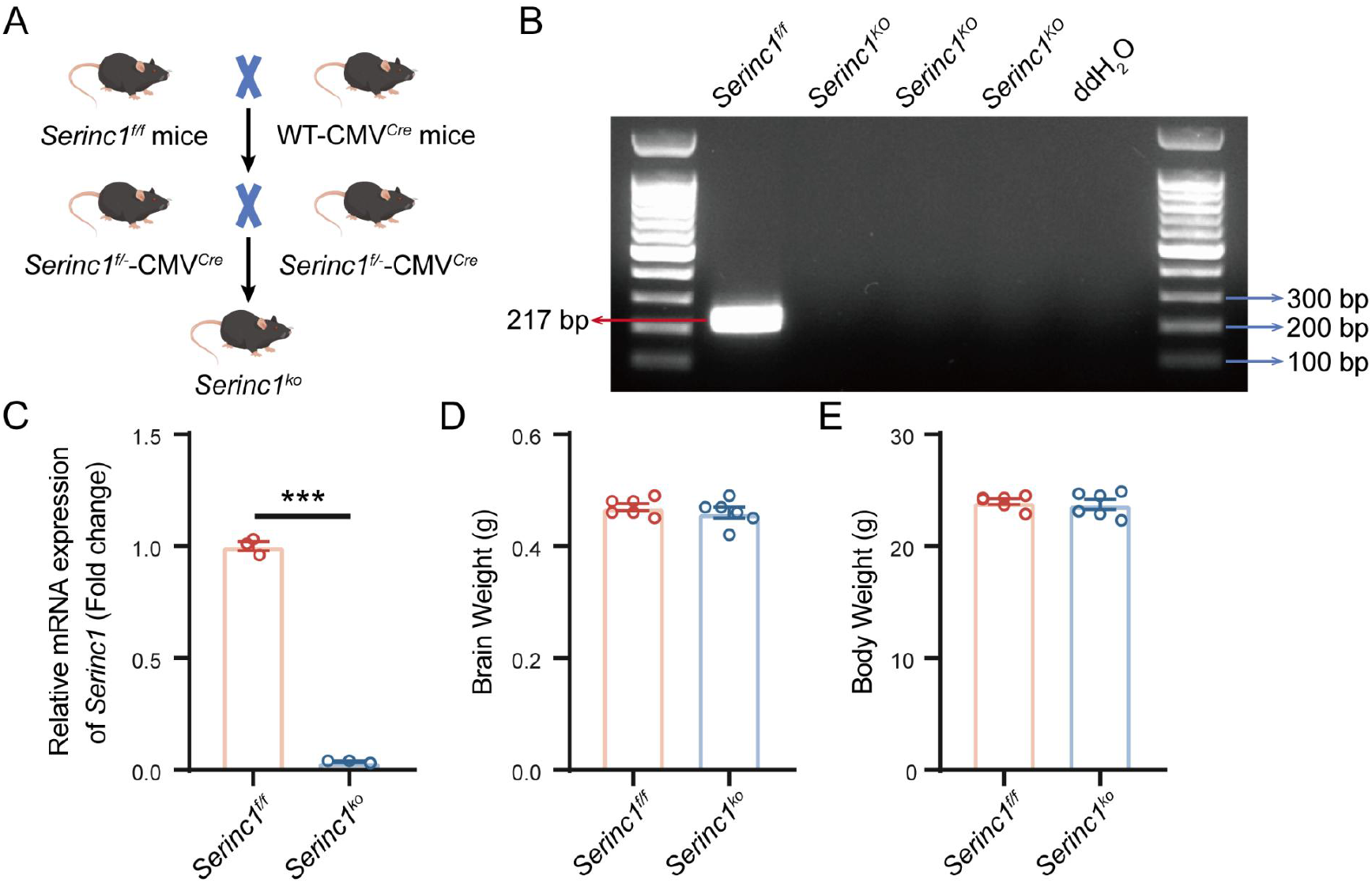
Generation and validation of the constitutive *Serinc1* knockout mouse model. **(A)** Schematic illustrating the generation of constitutive *Serinc1* knockout (KO) mice. *Serinc1^f/f^* mice were crossed with CMV-Cre transgenic mice to generate germline *Serinc1^ko^* mice. Intercrossing of the offspring yielded homozygous *Serinc1^ko^* mice for subsequent experiments. **(B)** Representative PCR genotyping of *Serinc1^ko^* mice. A recombination-specific PCR product (217 bp) was detected in genomic DNA from *Serinc1^ko^* mice, confirming successful Cre-mediated deletion of the *Serinc1* locus. **(C)** Validation of *Serinc1* deletion by quantitative RT–PCR. *Serinc1* mRNA expression was nearly abolished in *Serinc1^ko^* mice compared with *Serinc1^f/f^* littermate controls. **(D** and **E)** Quantification of brain weight (D) and body weight (E) in adult *Serinc1^f/f^* and *Serinc1^ko^* mice.

**Fig. S4.**
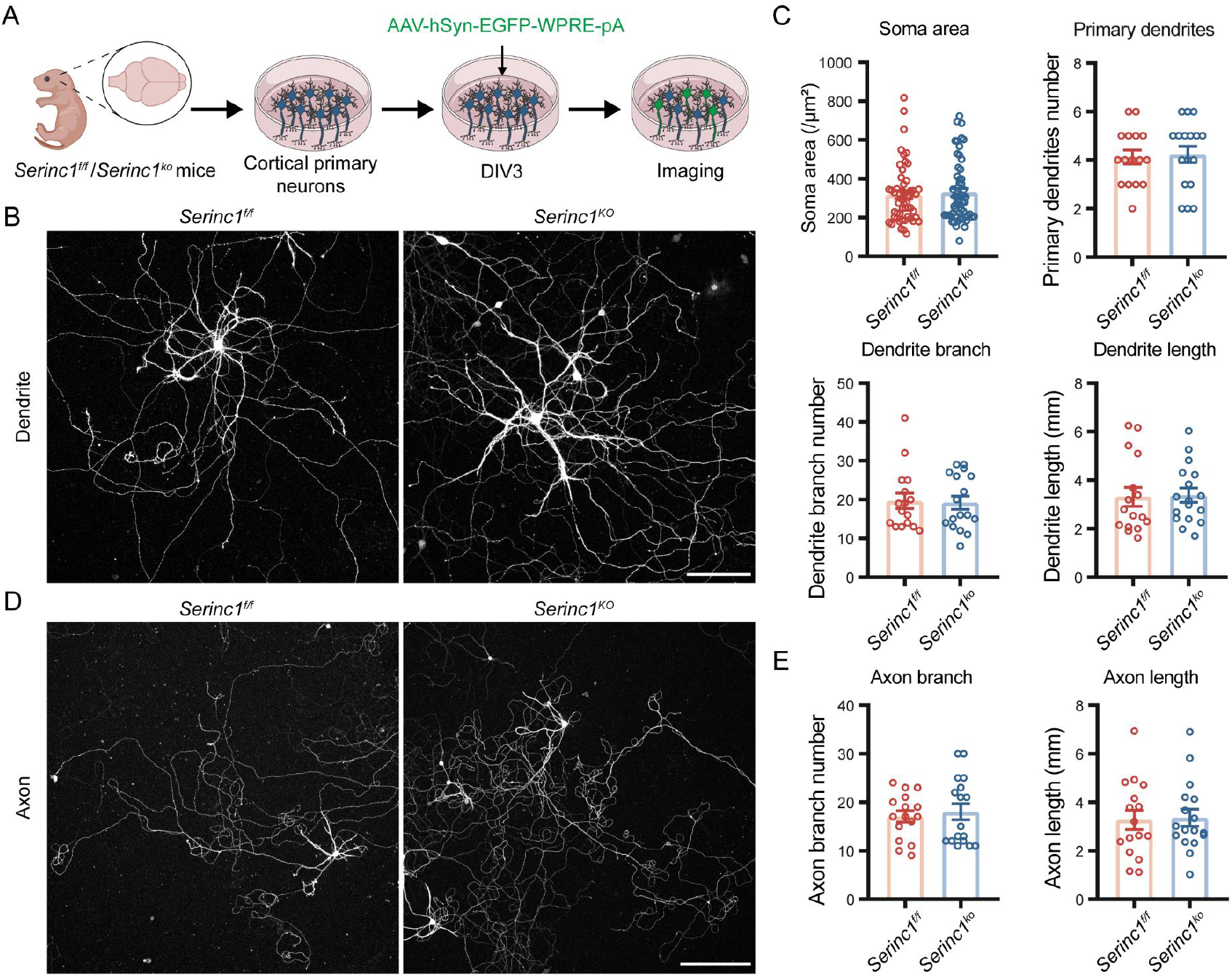
Loss of SERINC1 does not affect the morphological features of primary cortical neurons. **(A)** Schematic of the experimental design. Primary cortical neurons were isolated from *Serinc1^f/f^* and *Serinc1^ko^* mice, transduced with AAV-hSyn-EGFP at DIV3, and imaged at DIV14 for morphological analysis. **(B)** Representative fluorescence images of dendritic morphology in *Serinc1^f/f^* and *Serinc1^ko^* primary cortical neurons. Scale bar, 100 μm. **(C)** Quantification of neuronal soma area, the number of primary dendrites, the number of dendritic branches, and total dendritic length. **(D)** Representative fluorescence images of axonal morphology in *Serinc1^f/f^* and *Serinc1^ko^* primary cortical neurons. Scale bar, 200 μm. **(E)** Quantification of the number of axonal branches and total axonal length.

**Fig. S5.**
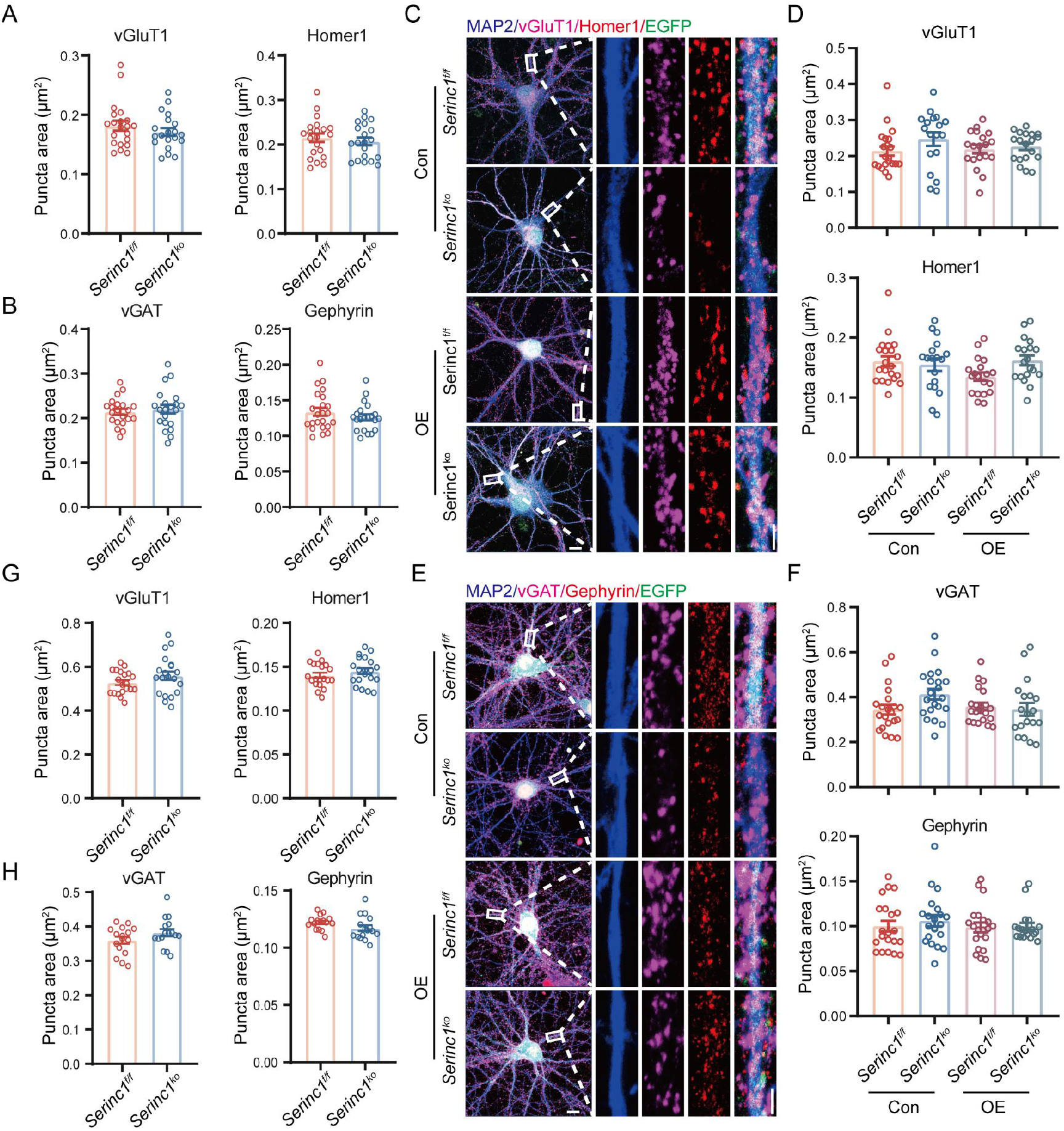
SERINC1 does not regulate the size of synaptic puncta in cortical neurons. **(A** and **B)** Quantification of excitatory synaptic puncta size labeled by the presynaptic marker vGluT1 and the postsynaptic marker Homer1 (A) and inhibitory synaptic puncta size labeled by the presynaptic marker vGAT and the postsynaptic marker Gephyrin (B) in primary cortical neurons derived from *Serinc1^f/f^* and *Serinc1^ko^* mice. **(C** and **D)** Representative immunofluorescence images (C) and quantification of excitatory synaptic puncta size (D) labeled by vGluT1 and Homer1 in primary cortical neurons derived from *Serinc1^f/f^* and *Serinc1^ko^* mice transduced at DIV3 with AAVs expressing EGFP (Con) or EGFP-P2A-SERINC1(OE) and analyzed at DIV14. Scale bars, 10 μm (insets, 2 μm). **(E** and **F)** Representative immunofluorescence images (E) and quantification of inhibitory synaptic puncta size (F) labeled by vGAT and Gephyrin in primary cortical neurons derived from *Serinc1^f/f^* and *Serinc1^ko^* mice transduced with control or *Serinc1*-overexpressing AAVs. Scale bars, 10 μm (insets, 2 μm). **(G** and **H)** Quantification of excitatory synaptic puncta size labeled by vGluT1 and Homer1 (G) and inhibitory synaptic puncta size labeled by vGAT and Gephyrin (H) in the ACC of adult *Serinc1^f/f^* and *Serinc1^ko^* mice.

**Fig. S6.**
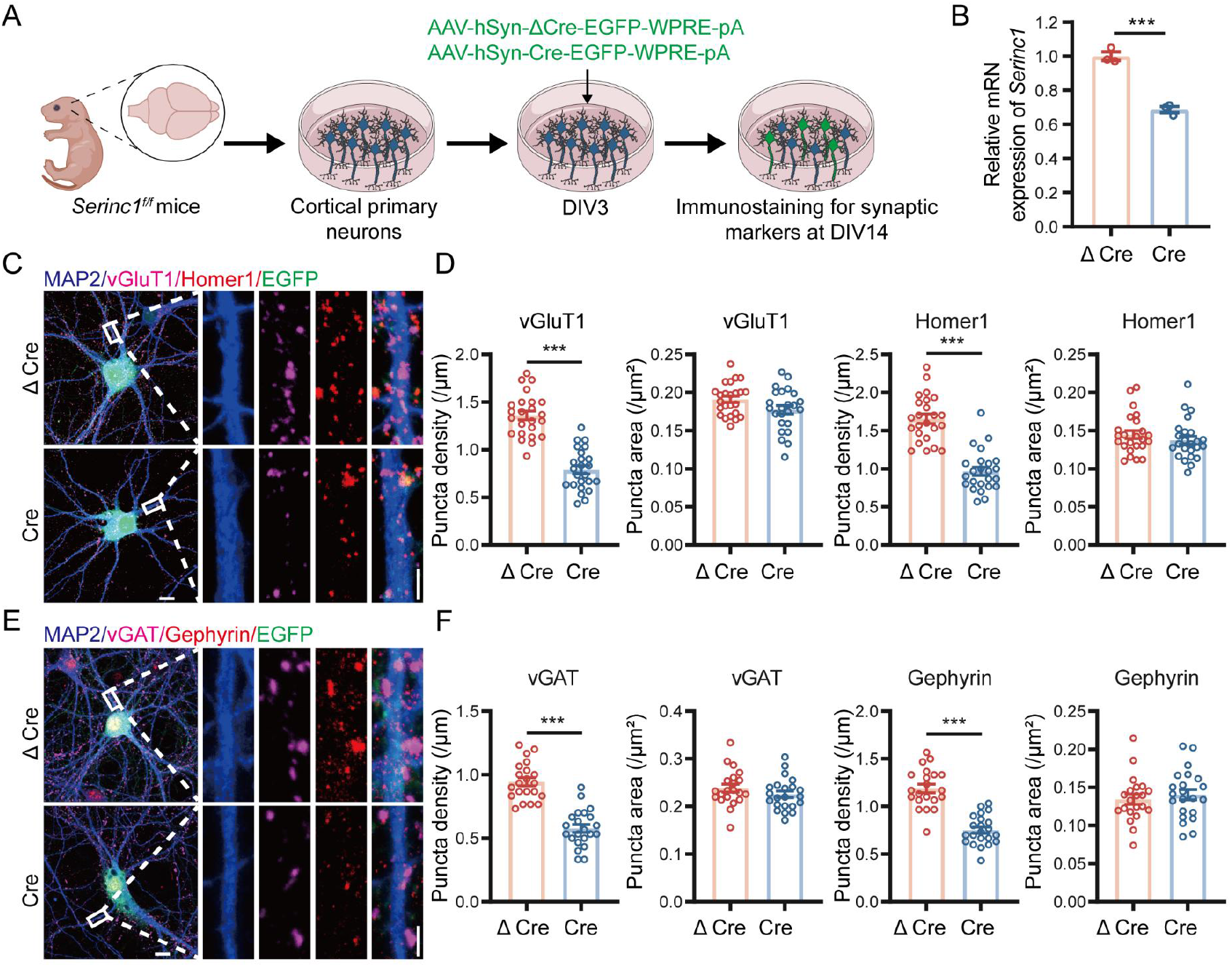
Neuron-specific deletion of *Serinc1* reduces synaptic puncta density in primary cortical neurons. **(A)** Schematic of the experimental design. Primary cortical neurons were isolated from *Serinc1^f/f^* mice, transduced at day in DIV3 with Cre- or ΔCre-expressing AAVs, and analyzed at DIV14 following immunofluorescence staining for synaptic markers. **(B)** Validation of *Serinc1* deletion by quantitative RT–PCR. Relative *Serinc1* mRNA expression in primary cortical neurons transduced with Cre- or ΔCre-expressing AAVs. **(C** and **D)** Representative immunofluorescence images (C) and quantification (D) of excitatory synaptic puncta density and size labeled by vGluT1 and Homer1 in ΔCre- and Cre-transduced primary cortical neurons. Scale bars, 10 μm (insets, 2 μm). **(E** and **F)** Representative immunofluorescence images (E) and quantification (F) of inhibitory synaptic puncta density and size labeled by vGAT and Gephyrin in ΔCre- and Cre-transduced neurons. Scale bars, 10 μm (insets, 2 μm).

**Fig. S7.**
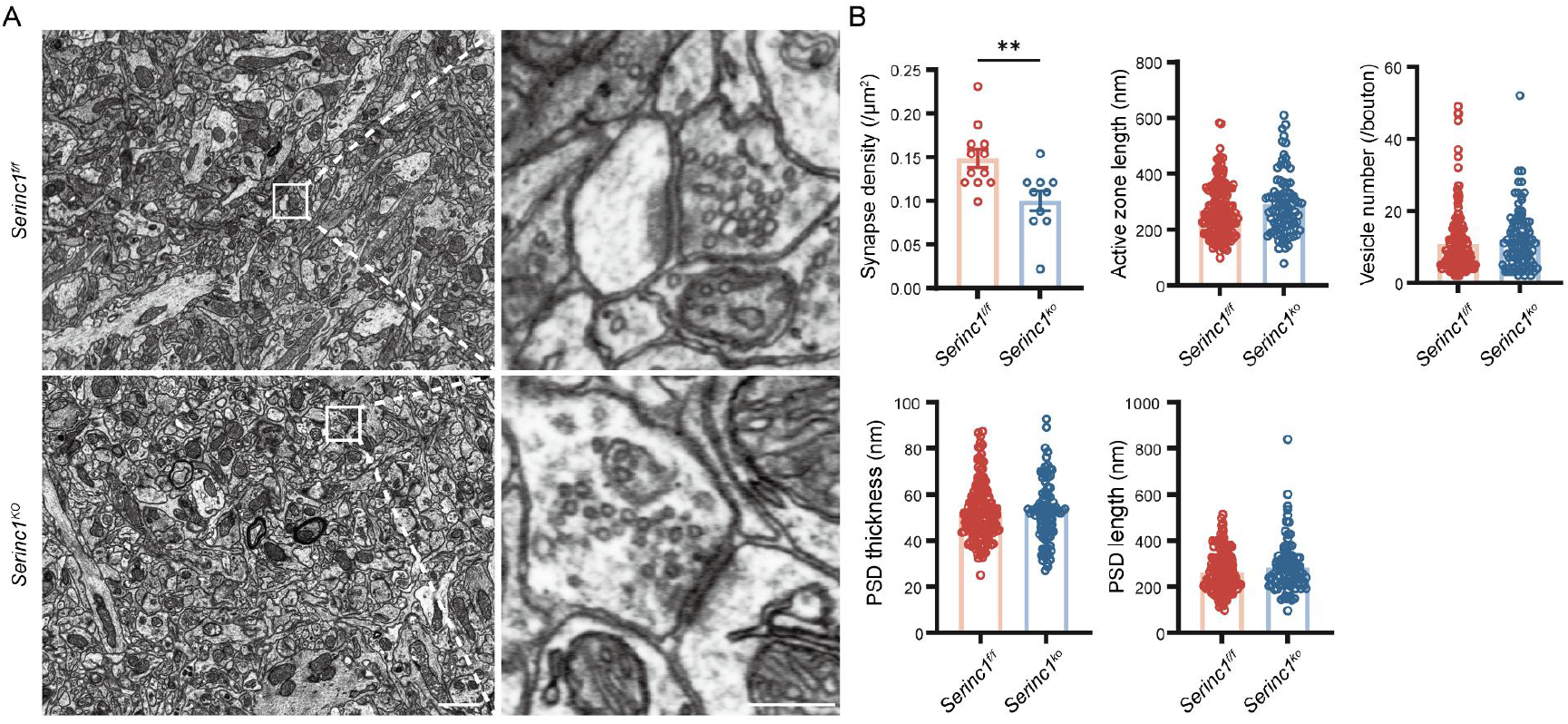
SERINC1 is required for maintaining synapse density but not synaptic ultrastructure in the ACC. **(A)** Representative transmission electron micrographs of Gray type I (asymmetric) excitatory synapses in the ACC of *Serinc1^f/f^* and *Serinc1^ko^* mice. Scale bars, 1 μm (insets, 200 nm). **(B)** Quantification of ultrastructural parameters of asymmetric excitatory synapses, including synapse density (synapses per μm²), active zone length, synaptic vesicle number per presynaptic terminal, PSD thickness, and PSD length, in the ACC of *Serinc1^f/f^* and *Serinc1^ko^* mice.

**Fig. S8.**
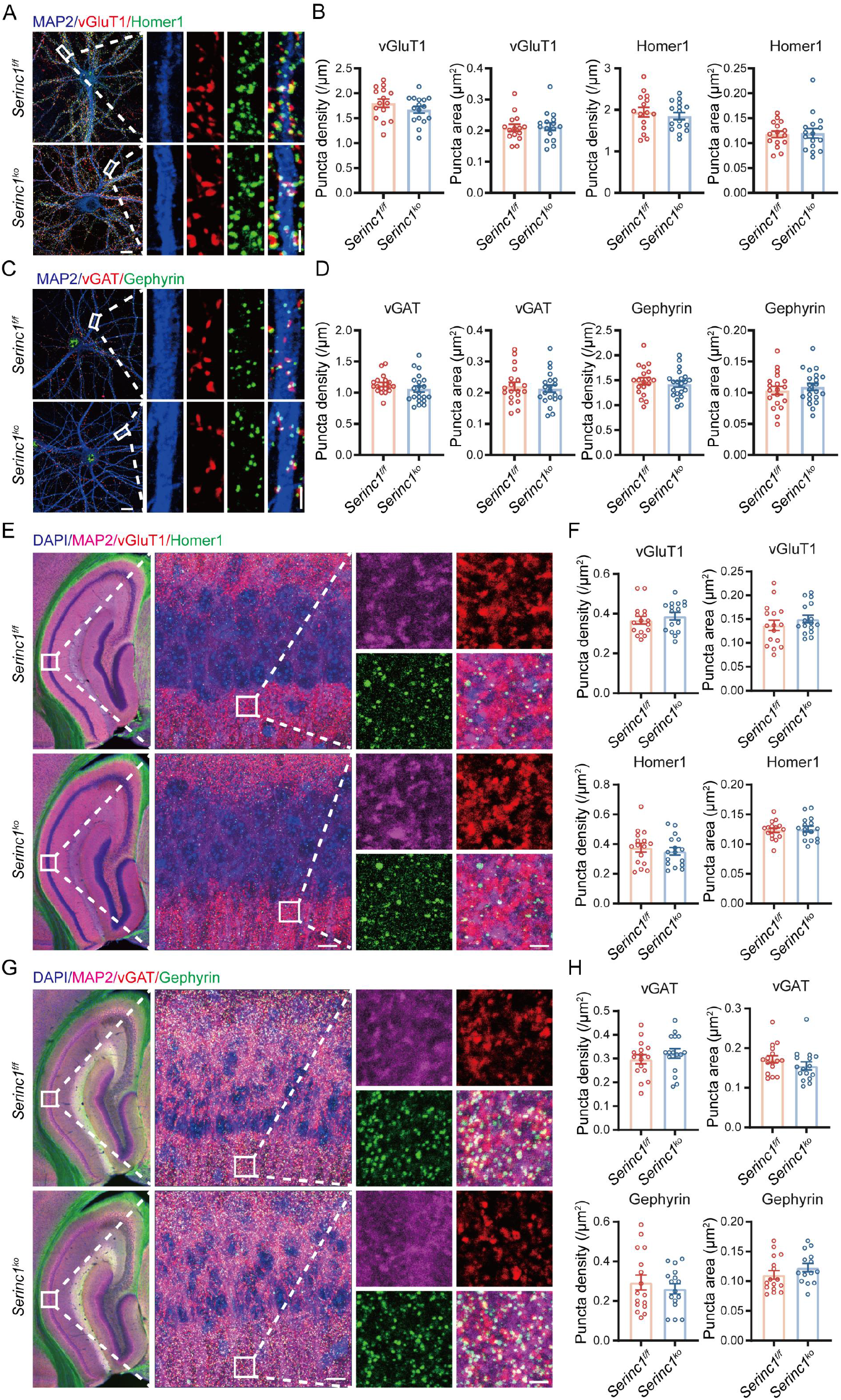
Loss of SERINC1 does not affect synapse density in hippocampal neurons. **(A** and **B)** Representative images (A) and quantification (B) of excitatory synaptic puncta labeled with the presynaptic marker vGluT1 and postsynaptic marker Homer1 in primary hippocampal neurons derived from *Serinc1^f/f^* and *Serinc1^ko^* mice. Scale bars, 10 μm (insets, 2 μm). **(C** and **D)** Representative images (C) and quantification (D) of inhibitory synaptic puncta labeled with the presynaptic marker vGAT and postsynaptic marker Gephyrin in primary hippocampal cortical neurons derived from *Serinc1^f/f^* and *Serinc1^ko^* mice. Scale bars, 10 μm (2 μm for zoomed in images). **(E** and **F)** Representative images (E) and quantification (F) of excitatory synaptic puncta labeled with the presynaptic marker vGluT1 and postsynaptic marker Homer1 in the hippocampal CA1 of adult *Serinc1^f/f^* and *Serinc1^ko^* mice. Scale bars, 10 μm (2 μm for zoomed in images). **(G** and **H)** Representative images (G) and quantification (H) of inhibitory synaptic puncta labeled with the presynaptic marker vGAT and postsynaptic marker Gephyrin in the hippocampal CA1 of adult *Serinc1^f/f^* and *Serinc1^ko^* mice. Scale bars, 10 μm (insets, 2 μm).

**Fig. S9.**
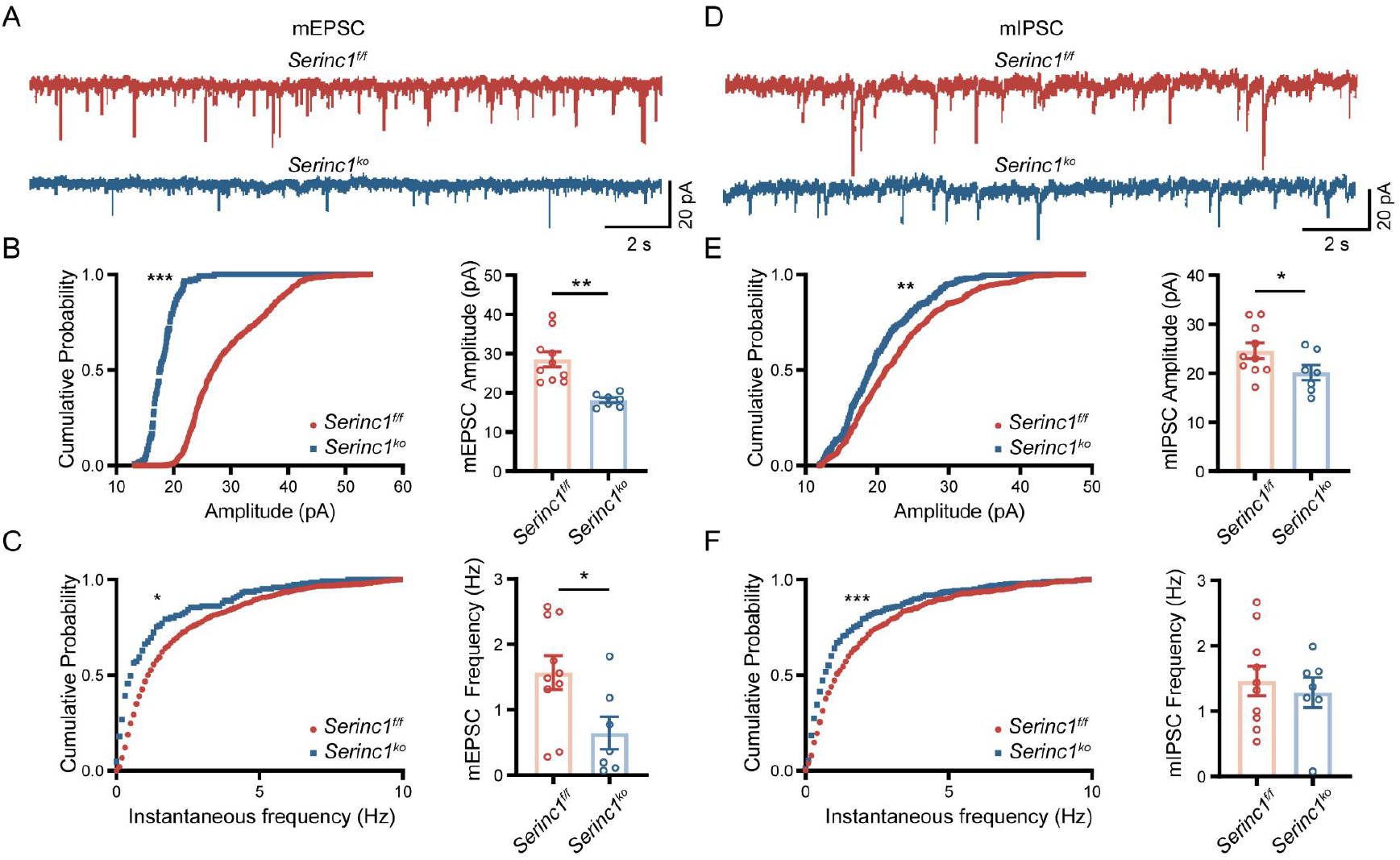
Loss of SERINC1 impairs synaptic transmission in primary cortical neurons. **(A)** Representative traces of mEPSCs recorded from primary cortical neurons derived from *Serinc1^f/f^* and *Serinc1^ko^* mice. **(B)** Cumulative probability distributions (left) and summary quantification (right) of mEPSC amplitude. **(C)** Cumulative probability distributions (left) and summary quantification (right) of mEPSC frequency. **(D)** Representative traces of miniature inhibitory postsynaptic currents (mIPSCs) recorded from primary cortical neurons derived from *Serinc1^f/f^* and *Serinc1^ko^* mice. **(E)** Cumulative probability distributions (left) and summary quantification (right) of mIPSC amplitude. **(F)** Cumulative probability distributions (left) and summary quantification (right) of mIPSC frequency.

**Fig. S10.**
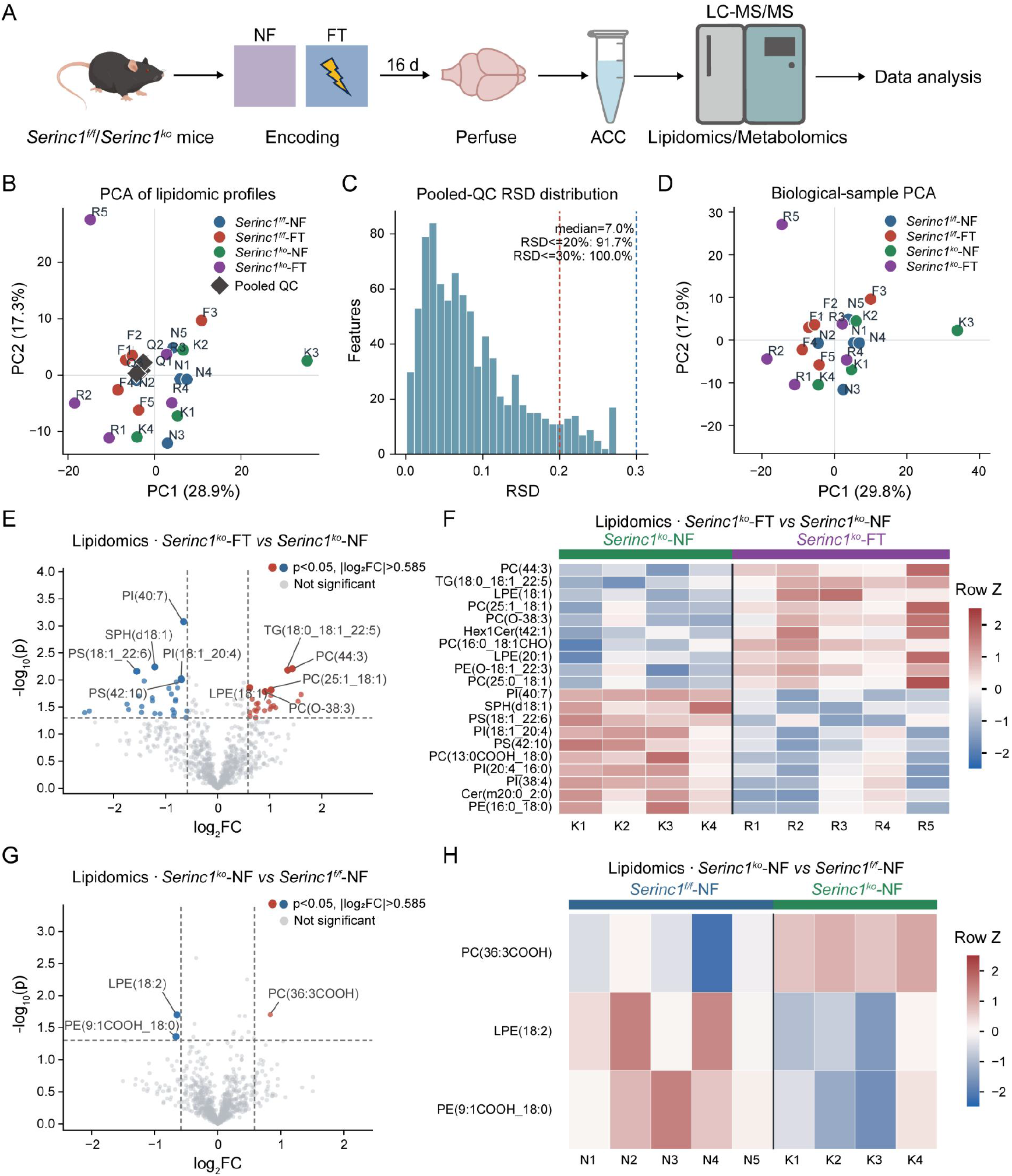
Lipidomic profiling of the ACC in *Serinc1*-deficient mice following fear conditioning. **(A)** Schematic illustration of the experimental design for lipidomic and metabolomic profiling of the ACC in *Serinc1^f/f^* and *Serinc1^ko^* mice. Mice underwent encoding and were subjected to tissue collection 16 days later. ACC tissues were isolated and analyzed by liquid chromatography–tandem mass spectrometry (LC-MS/MS). **(B)** Principal component analysis (PCA) of lipidomic profiles from *Serinc1^f/f^* and *Serinc1^ko^* mice under NF and FT conditions. Each point represents an individual biological sample; pooled quality-control (QC) samples are indicated by black diamonds. **(C)** Distribution of relative standard deviations (RSDs) across features in pooled QC samples. The median RSD was 7.0%, with 91.7% and 100% of detected features exhibiting RSDs below 20% and 30%, respectively. **(D)** PCA of biological samples based on lipidomic profiles. Each point represents an individual biological sample, with sample identities indicated by labels. **(E)** Volcano plot showing differentially abundant lipid species between *Serinc1^ko^*-FT and *Serinc1^ko^*-NF groups. Lipid species meeting the indicated statistical and fold-change criteria are highlighted. **(F)** Heatmap showing the relative abundance of significantly altered lipid species between *Serinc1^ko^*-FT and *Serinc1^ko^*-NF groups. Each column represents an individual biological sample, and lipid abundance is displayed as row-wise Z scores. **(G)** Volcano plot showing differentially abundant lipid species between *Serinc1^ko^*-NF and *Serinc1^f/f^*-NF groups. Lipid species meeting the indicated statistical and fold-change criteria are highlighted. **(H)** Heatmap showing the relative abundance of significantly altered lipid species between *Serinc1^ko^*-NF and *Serinc1^f/f^*-NF groups. Each column represents an individual biological sample, and lipid abundance is displayed as row-wise Z scores.

**Fig. S11.**
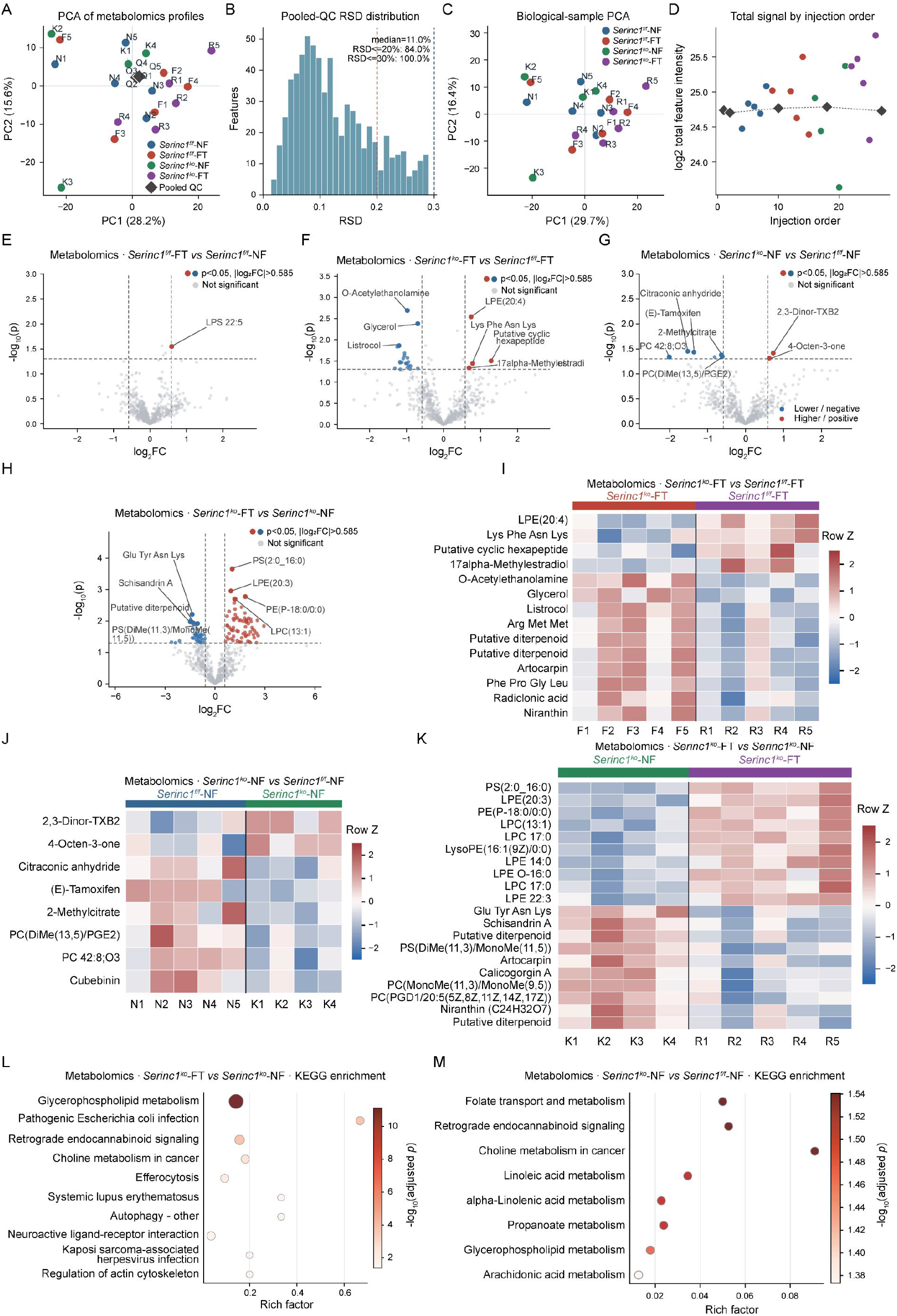
Metabolomic profiling of the ACC in *Serinc1*-deficient mice following fear conditioning. **(A)** PCA of metabolomic profiles from the ACC of *Serinc1^f/f^* and *Serinc1^ko^* mice under NF and FT conditions. Each point represents an individual biological sample, and pooled QC samples are indicated by black diamonds. **(B)** Distribution of relative standard deviations (RSDs) across detected metabolites in pooled QC samples. **(C)** PCA of biological samples based on metabolomic profiles. Each point represents an individual biological sample. **(D)** Total metabolite signal intensity plotted against injection order to assess signal stability throughout the LC-MS/MS acquisition sequence. **(E)**Volcano plot showing differentially abundant metabolites between *Serinc1^f/f^*-FT and *Serinc1^f/f^*-NF groups. Metabolites meeting the criteria of P < 0.05 and |log₂FC| > 0.585 are highlighted. **(F)** Volcano plot showing differentially abundant metabolites between *Serinc1^ko^*-FT and *Serinc1^f/f^*-FT groups. Metabolites meeting the criteria of P < 0.05 and |log₂FC| > 0.585 are highlighted. **(G)** Volcano plot showing differentially abundant metabolites between *Serinc1^ko^*-NF and *Serinc1^f/f^*-NF groups. Metabolites meeting the criteria of P < 0.05 and |log₂FC| > 0.585 are highlighted. **(H)** Volcano plot showing differentially abundant metabolites between *Serinc1^ko^*-FT and *Serinc1^ko^*-NF groups. Metabolites meeting the criteria of P < 0.05 and |log₂FC| > 0.585 are highlighted. **(I)** Heatmap showing the relative abundance of differentially abundant metabolites between *Serinc1^ko^*-FT and *Serinc1^f/f^*-FT groups. Each column represents an individual biological sample, and metabolite abundance is displayed as row-wise Z scores. **(J)** Heatmap showing the relative abundance of differentially abundant metabolites between *Serinc1^ko^*-NF and *Serinc1^f/f^*-NF groups. Each column represents an individual biological sample, and metabolite abundance is displayed as row-wise Z scores. **(K)** Heatmap showing the relative abundance of differentially abundant metabolites between *Serinc1^ko^*-FT and *Serinc1^ko^*-NF groups. Each column represents an individual biological sample, and metabolite abundance is displayed as row-wise Z scores. **(L)** KEGG pathway enrichment analysis of differentially abundant metabolites between *Serinc1^f/f^*-FT and *Serinc1^f/f^*-NF groups. The x axis indicates the rich factor, and dot color indicates the statistical significance of pathway enrichment. **(M)** KEGG pathway enrichment analysis of differentially abundant metabolites between *Serinc1^ko^*-FT and *Serinc1^ko^*-NF groups. The x axis indicates the rich factor, and dot color indicates the statistical significance of pathway enrichment.

**Fig. S12.**
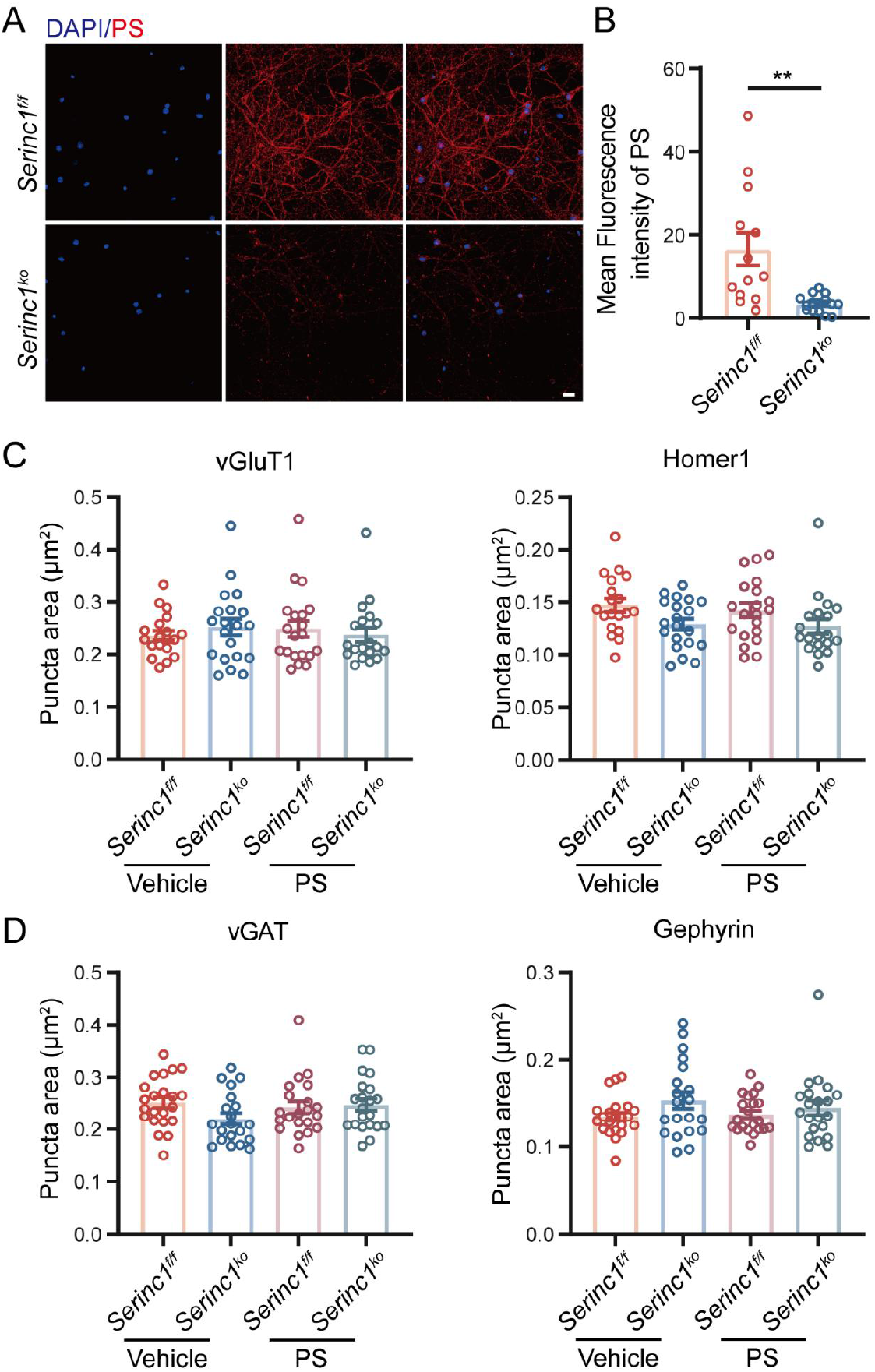
Loss of SERINC1 reduces Annexin V-detected PS signal in primary cortical neurons. **(A** and **B)** Representative images (A) and quantification (B) of Annexin V–mCherry-detected PS signal in 4% PFA-fixed primary cortical neurons derived from *Serinc1^f/f^* and *Serinc1^ko^* mice. Scale bar, 20 μm. **(C** and **D)** Quantification of excitatory synaptic puncta size labeled by the presynaptic marker vGluT1 and the postsynaptic marker Homer1 (C) and inhibitory synaptic puncta size labeled by the presynaptic marker vGAT and the postsynaptic marker Gephyrin (D) in primary cortical neurons derived from *Serinc1^f/f^* and *Serinc1^ko^* mice following treatment with vehicle or PS (20 μg ml⁻¹) for 48 h beginning at DIV12.

**Fig. S13.**
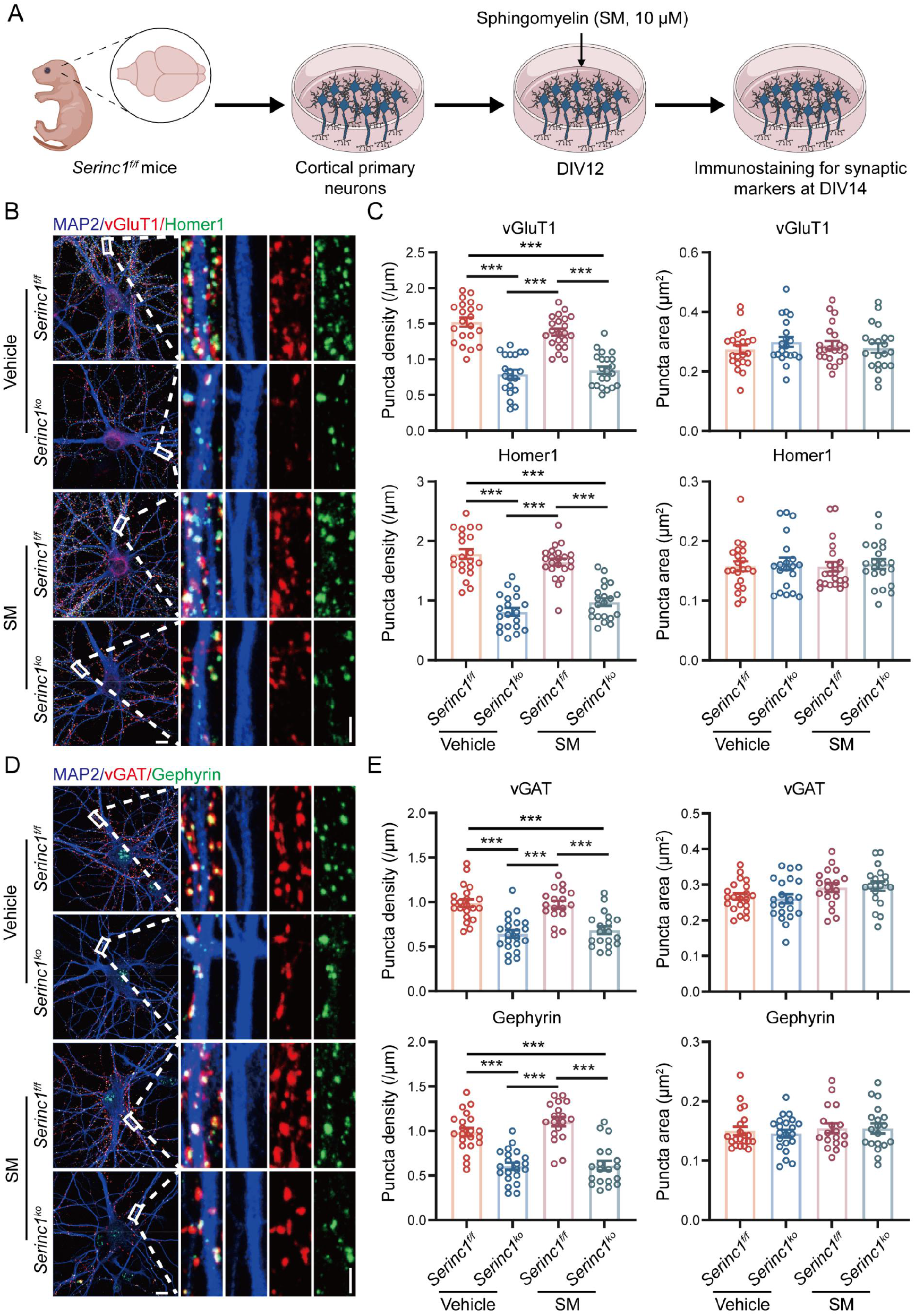
Sphingomyelin supplementation fails to rescue the reduction of synaptic puncta density caused by SERINC1 deficiency in primary cortical neurons. **(A)** Schematic of the experimental design. Primary cortical neurons were isolated from *Serinc1^f/f^* and *Serinc1^ko^* mice, treated at day in DIV12 with vehicle or sphingomyelin (SM; 10 μM), and analyzed at DIV14 following immunofluorescence staining for synaptic markers. **(B** and **C)** Representative immunofluorescence images (B) and quantification (C) of excitatory synaptic puncta density and size labeled by vGluT1 and Homer1 in primary cortical neurons derived from *Serinc1^f/f^* and *Serinc1^ko^* mice following vehicle or SM treatment. Scale bars, 10 μm (insets, 2 μm). **(D** and **E)** Representative immunofluorescence images (D) and quantification (E) of inhibitory synaptic puncta density and size labeled by vGAT and Gephyrin in primary cortical neurons derived from *Serinc1^f/f^* and *Serinc1^ko^* mice following vehicle or SM treatment. Scale bars, 10 μm (insets, 2 μm).

